# Endogenous retroviruses drive KRAB zinc-finger family protein expression for tumor suppression

**DOI:** 10.1101/2020.02.02.931501

**Authors:** Jumpei Ito, Izumi Kimura, Andrew Soper, Alexandre Coudray, Yoshio Koyanagi, Hirofumi Nakaoka, Ituro Inoue, Priscilla Turelli, Didier Trono, Kei Sato

## Abstract

Numerous genes are aberrantly expressed in tumors, but its cause remains unclear. Human endogenous retroviruses (HERVs) are repetitive elements in the genome and have a potential to work as enhancers modulating adjacent genes. Since numerous HERVs are activated epigenetically in tumors, their activation could alter gene expression globally in tumors and change the tumor characteristics. Here, we show the HERV activation in tumors is associated with the upregulation of hundreds of transcriptional suppressors, Krüppel-associated box domain-containing zinc-finger family proteins (KZFPs). KZFP genes are preferentially encoded nearby the activated HERVs in tumors and transcriptionally regulated by the adjacent HERVs. Increased HERV and KZFP expression in tumors was associated with better disease conditions. Many KZFPs could suppress the progressive characteristics of cancer cells by downregulating genes related to the cell cycle and cell-matrix adhesion. Our data suggest that HERV activation in tumors drives the concerted expression of KZFP genes for tumor suppression.

## Introduction

Aberrant gene expression is a hallmark of cancers. Gene expression statuses in tumors are highly diverse among patients and are associated with the phenotypes of tumors such as proliferation, invasion/metastasis capacity, and therapeutic response as well as the clinical outcome of patients^1^. Particularly, many genes that are aberrantly expressed in tumors and associated with cancer progression have been identified^2^; however, the abnormality of the gene regulatory network underlying the aberrant expression of these genes in tumors is poorly understood^3–5^.

Decades of research have highlighted the significance of regulatory sequences derived from human endogenous retroviruses (HERVs) in the modulation of human gene expression^6^. HERVs are a type of transposable element (TE) that originates from ancient retroviral infection in host germ cells^7^. There are several hundred types of HERVs in the human genome, constituting 8% of the genome^8^. Unlike other TEs, HERVs possess long terminal repeat (LTR) sequences that particularly densely contain transcriptional regulatory elements^9,10^ and function as viral promoters^7^. In addition, HERV LTRs have the potential to function as promoters or enhancers of adjacent genes^6^. While most HERVs are epigenetically silenced in normal tissues, some HERVs function as part of the host gene regulatory network and play crucial roles in diverse biological events^6,11–16^. For instance, HERVs harboring STAT1- and IRF1-binding sites are essential for the interferon inducibility of genes related to the innate immune response^17^.

The expression of HERVs in normal tissues is controlled by epigenetic mechanisms such as DNA methylation and repressive histone modifications^18,19^; in contrast, HERV expression is highly elevated in various types of cancers^20–24^. Since the elevation of HERV expression in tumors is presumably caused by epigenetic reactivation, the expressed HERVs could upregulate the expression of adjacent genes. Therefore, it is possible that the derepression of numerous HERVs in tumors globally alters host gene expression and changes the characteristics of cancers^25,26^. To test this hypothesis, we investigated the multi-omics dataset of tumors provided by The Cancer Genome Atlas (TCGA)^27^ and assessed the effects of HERV activation on host gene expression. We found that genome-wide HERV activation in tumors is associated with the upregulation of potent transcriptional suppressor genes, Krüppel-associated box (KRAB) domain-containing zinc-finger family protein (KZFP) genes^28^, which are preferentially located in the vicinity of activated HERVs. Although KZFPs are widely known as transcriptional silencers against TEs, including HERVs^28^, our data highlight that the expression of KZFP genes is induced by the adjacent HERVs in tumors, leading to global gene expression alterations and phenotypic changes.

## Results

### Characterization of expressed HERVs across 12 types of solid tumors

We investigated the tumor RNA-sequencing (RNA-Seq) data of 5,470 patients provided by TCGA (**Data S1**). Only RNA-Seq reads that were uniquely mapped to the human genome were analyzed. A total of 11,011 loci of expressed HERVs were identified across twelve types of solid tumors (**Fig. 1A and Data S2**). While some HERVs were detected in only specific types of cancers, the majority of the expressed HERVs were detected in multiple types of cancers, and the sets of the expressed HERV loci were highly similar among all cancer types (**Figs. S1A and S1B**). In nine out of the twelve types of cancers, the overall expression levels of HERVs were increased compared to that in the adjacent normal tissues (**Fig. 1B**), consistent with previous reports^20–24^. Dimension reduction analysis based on HERV expression profiles showed that each type of cancer displays a distinguishable pattern of HERV expression (**Fig. 1C**). Importantly, the expressed HERVs preferentially overlapped with the nucleosome-free regions (NFRs) determined by Assay for Transposase-Accessible Chromatin Sequencing (ATAC-Seq) (**Fig. 1D**), suggesting that expressed HERVs in tumors are epigenetically active and have the potential to modulate adjacent gene expression.

**Fig. 1.**
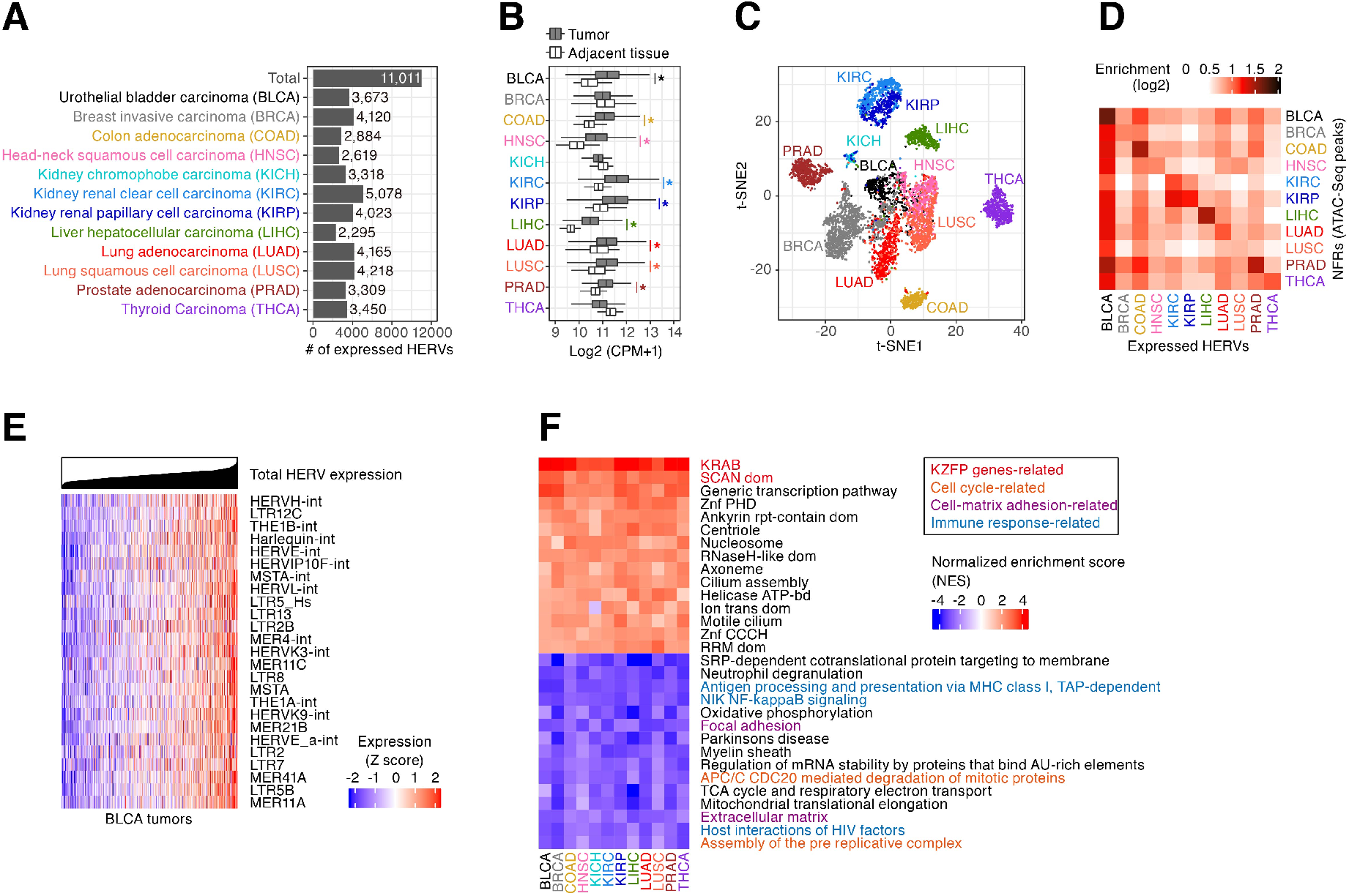
Landscape of HERV expression in 12 types of solid cancers. A) Numbers of the expressed HERV loci identified in respective types of cancers. B) Total expression levels of HERVs (log2 (counts per million (CPM) + 1)) in cancers and adjacent normal tissues. An asterisk denotes a significant increase in the values in tumors compared to that in normal tissues (Bonferroni-corrected *P* value < 0.05 in two-sided Wilcoxon rank sum test). C) t-SNE plot representing the expression patterns of HERVs among tumor samples. Dots indicate tumor sample data. The expression levels of the 1000 most highly expressed HERVs were used in the analysis. D) Fold enrichments of the overlaps between expressed HERV loci and nucleosome-free regions (NFRs; i.e., ATAC-Seq peaks) identified in respective types of cancers. The enrichment value was calculated based on the random expectation. E) Expression levels of the respective HERV groups in BLCA tumors. Normalized expression levels (Z scores) of the 25 most highly expressed HERV groups are shown. Tumors were ordered according to the total value. F) Gene set enrichment analysis (GSEA)^29^ summarizing genes whose expression levels were correlated with the global expression levels of HERVs. Spearman’s correlation scores between the expression levels of respective genes and the total expression level of HERVs were calculated, and GSEA was subsequently performed based on those scores. For the positive (red) and negative (blue) correlations, the high-scored 15 gene sets (regarding the mean value among cancer types) are shown. Redundant gene sets were removed from the results.

### Transcriptome signatures associated with the global derepression of HERVs in tumors

Although HERV expression levels tended to be elevated in tumors compared to the corresponding normal tissues (**Fig. 1B**), the genome-wide expression levels of HERVs in tumors were highly heterogeneous among patients, even within the same cancer type (**Figs. S1C and 1E**). Notably, such global HERV activation occurred regardless of the type of HERV (**Figs. 1E, S1D, and S1E**), although the regulatory sequences of these HERVs were highly diverse^10^. In many types of cancers, the global expression levels of HERVs were negatively correlated with the DNA methylation levels of CpG sites that are on or proximal (<1 kb) to the expressed HERVs (**Figs. S1F and S1G**), suggesting that the epigenetic derepression of HERVs is a cause of the elevation of HERV expression in tumors.

To elucidate the effects of the global derepression of HERVs on host gene expression in tumors, we investigated the genes whose expression was associated with HERV derepression in tumors. We assessed the correlation of the expression level of each gene with the total expression level of HERVs in tumors and subsequently performed gene set enrichment analysis (GSEA)^29^ based on the above correlation scores. We found that the genes showing a correlation with HERVs were highly similar among distinct types of cancers (**Figs. S2A and S2B**). KZFP genes (i.e., genes possessing the KRAB domain) were highly upregulated upon the elevation of HERV expression (**Fig. 1F**). Most KZFP genes were co-expressed with each other (**Fig. S2C**) and with the major groups of HERVs in tumors (**Fig. S2D**). Additionally, genes related to the cell cycle, cell-matrix adhesion, and immune response were downregulated upon the upregulation of HERV and KZFP genes (**Figs. 1F and S3**). We investigated another RNA-Seq dataset of cancer cell lines provided by the Cancer Cell Line Encyclopedia (CCLE)^30^ and verified that the expression of HERVs was positively associated with KZFP genes and negatively associated with genes related to the cell cycle, cell-matrix adhesion, and immune response (**Fig. S4**). These results suggest that these associations are arise from the expressional changes that occur in cancer cells themselves.

### Transcriptional activation of KZFP genes by surrounding HERVs

We hypothesized that derepressed HERVs near KZFP genes induce the expression of these genes, leading to the synchronized expression of HERVs and KZFP genes in tumors. It is known that KZFP genes form genomic clusters, particularly on chromosome 19 in the human genome^31^. We found that the expressed HERVs in tumors were preferentially present in these clusters of KZFP genes (**Fig. 2A**). The expressed HERVs in tumors and those with transcriptional regulatory signals (i.e., NFRs or enhancers defined by GeneHancer^32^) were highly enriched in the vicinity of transcriptional start sites (TSSs) of KZFP genes (**Figs. 2B and S5A**). Several types of HERV LTRs, such as LTR70, LTR25, LTR5B, and LTR5Hs, showed particularly strong enrichments around the TSSs of KZFP genes (**Fig. 2C**).

**Fig. 2.**
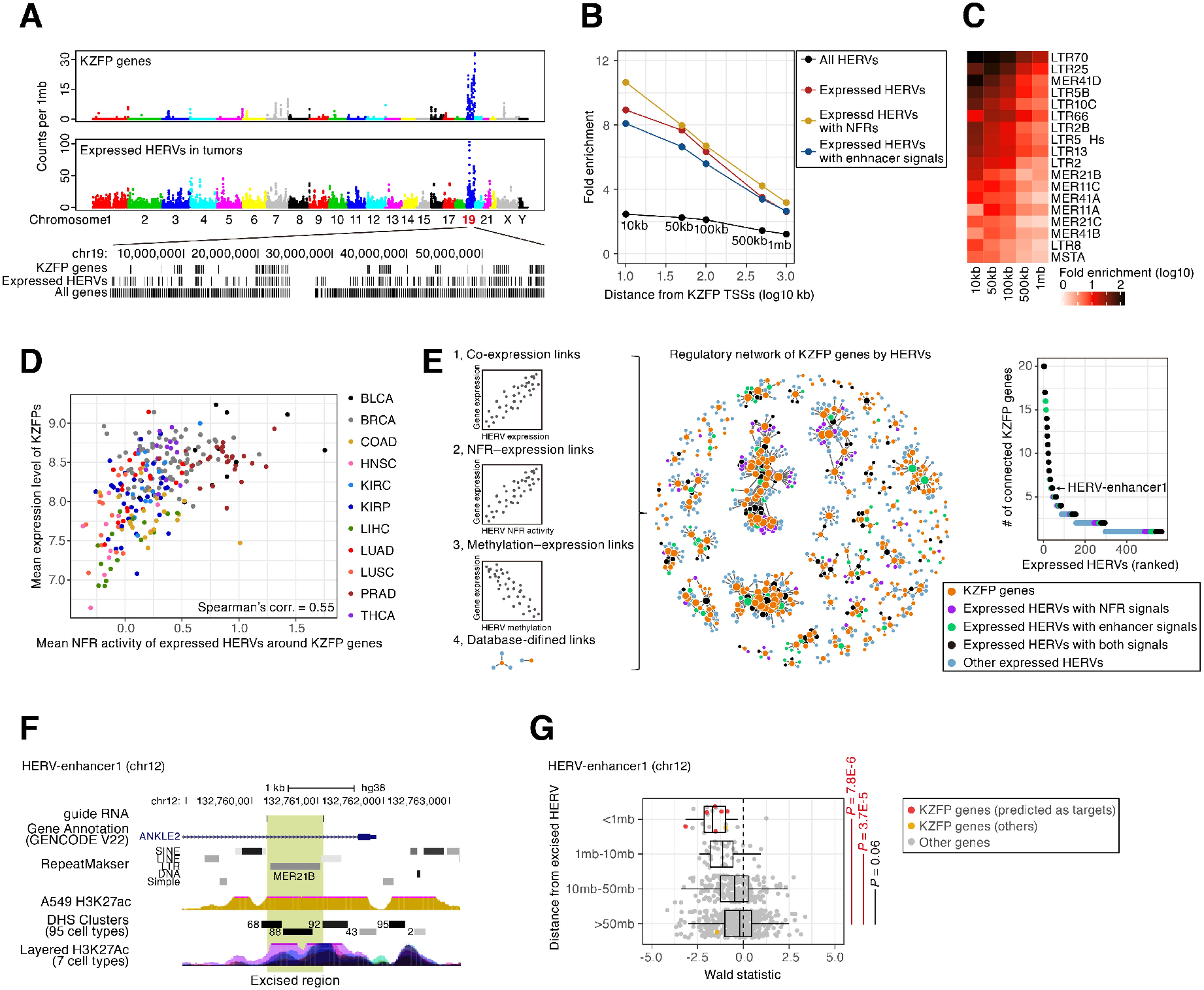
Transcriptional activation of KZFP genes by the adjacent HERVs. A) Genomic positions of KZFP genes and the expressed HERVs in tumors. Top) The genomic densities of KZFP genes and the expressed HERVs (counts per 1 megabase pairs (mb)). Bottom) Genomic locations of KZFP genes, the expressed HERVs, and all genes on chromosome 19. B) Enrichments of the expressed HERVs in tumors around the transcription start sites (TSSs) of KZFP genes. Fold enrichments of the four categories of HERVs (all HERVs, expressed HERVs, expressed HERVs with NFRs, and expressed HERVs overlapped with the enhancers defined by GeneHancer^32^) in the regions within 10, 50, 100, and 500 kb and 1 mb from the TSSs of KZFP genes are shown. The enrichment value was calculated based on the random expectation. C) Fold enrichments of respective groups of expressed HERVs (LTRs) in the vicinity of the TSSs of KZFP genes. LTR groups that were significantly (FDR < 0.05) enriched within 50 kb from the TSSs are shown. D) Association between the mean expression level of KZFPs and the mean NFR activity of the expressed HERVs in the vicinity (<50 kb) of KZFP genes in tumors. E) Prediction of the genes regulated by the expressed HERVs. Left) Schematics of the prediction of the regulatory relationships. The prediction was based on the following information: 1) co-expression interactions, 2) NFR–expression interactions, 3) DNA methylation–expression anti-correlation interactions, and 4) interactions predicted by GeneHancer^32^. The co-expression interaction was used in only pairs of HERV and KZFP genes within 50 kb of each other, while the other interactions were used in only pairs within 500 kb of each other. Middle) Integrated network representing the predicted regulations of KZFP genes by HERVs. Right) Numbers of connected KZFPs of the respective HERV nodes in the network. The HERVs were ranked according to connectivity. The target HERV for the CRISPR-Cas9 excision experiment is denoted. F) UCSC genome browser view of the target HERV (HERV-enhancer1). G) Effect of the excision of HERV on the expression of the adjacent genes in lung adenocarcinoma (A549) cells. The cells in which the target HERV was homozygously excised (5 clones) and the non-target control cells (5 clones) were compared. The X-axis indicates the Wald statistic, in which the positive and negative values indicate the up- and downregulation, respectively, of the gene expression compared to that in the non-target control cells. Genes were stratified according to the distance from the excised HERV, and the distributions of Wald statistics were compared between the indicated categories. *P* values were calculated by two-sided Student’s t-test.

We next investigated the association between the transcriptional upregulation of KZFP genes and the epigenetic activation of the adjacent HERVs in tumors. The mean expression level of KZFP genes was associated with the mean NFR activity of the expressed HERVs around those genes in tumors (**Fig. 2D**). Additionally, the mean expression level of KZFP genes in tumors was negatively correlated with the mean DNA methylation level of the CpG sites that are on or proximal (<1 kb) to the expressed HERVs around those genes (**Fig. S5B**). These findings suggest that the expression of KZFP genes in tumors is upregulated by the epigenetic derepression of adjacent HERVs.

Next, we searched for genes possibly regulated by respective HERV loci according to the co-expression, NFR–expression, and DNA methylation– expression relationships as well as the pre-defined enhancer–gene links^32^ (**Fig. 2E, left**) (**Data S3**). In these four types of predictions, KZFP genes were highly enriched in the set of genes possibly regulated by HERVs (**Fig. S5C**), supporting the significance of HERVs in the transcriptional regulation of KZFP genes. Based on these interactions, we constructed a network representing the regulation of KZFP genes by HERVs (**Fig. 2E, middle**). We identified several “hub” HERV loci, which are connected to many KZFP genes in the network and are likely to be involved in the transcriptional regulation of these genes (**Fig. 2E, right**).

To experimentally address the significance of HERVs in the transcriptional modulation of KZFP genes in cancer cells, we performed CRISPR-Cas9 excision of a hub HERV locus (HERV-enhancer1; **Fig. 2E, right**) in human lung adenocarcinoma (LUAD) (A549) cells (**Figs. 2F and S6**). We particularly selected this HERV locus because it displayed active histone marks in A549 cells (**Fig. 2F**). We demonstrated that the homozygous excision of this HERV decreased the expression of adjacent genes, including many KZFP genes (**Fig. 2G**).

### Biological relevance of the expression status of KZFPs and HERVs to cancer progression

Since KZFPs are potent transcriptional suppressors^28^, it is possible that the synchronized induction of many KZFPs in tumors would alter gene expression globally and change the characteristics of tumors. We found that somatic mutations accumulated particularly in the DNA-binding interfaces of KZFPs in tumors (**Fig. 3A**), suggesting that the aberration of the DNA-binding activity of KZFPs is associated with tumor progression. We therefore investigated the associations of the expression of KZFPs and HERVs with the clinical outcomes of cancer patients and found the following marked associations: in four (bladder carcinoma (BLCA), head and neck squamous cell carcinoma (HNSC), kidney renal papillary cell carcinoma (KIRP), and LUAD) out of twelve types of cancers, patients with high expression levels of KZFPs and HERVs in the tumors tended to show a better prognosis than those with low expression levels (**Figs. 3B and S7**). Furthermore, we examined the association of the expression levels of respective genes and HERV loci with cancer prognosis and found that KZFP genes and HERVs tended to show a stronger association with better prognosis than the other genes (**Figs. 3C, 3D**, and **S8**). Conversely, genes related to the cell cycle and cell-matrix adhesion tended to show a stronger association with a worse prognosis (**Fig. S8B**). We further examined the association of the overall expression level of KZFPs and cancer stage, which reflects the degree of invasion and metastasis of tumors. The overall expression level of KZFPs decreased as the cancer stage progressed in multiple types of cancers (**Figs. 3E, 3F, and S9**). Conversely, genes related to the cell cycle and cell-matrix adhesion increased as the cancer stage progressed (**Fig. S9B**).

**Fig. 3.**
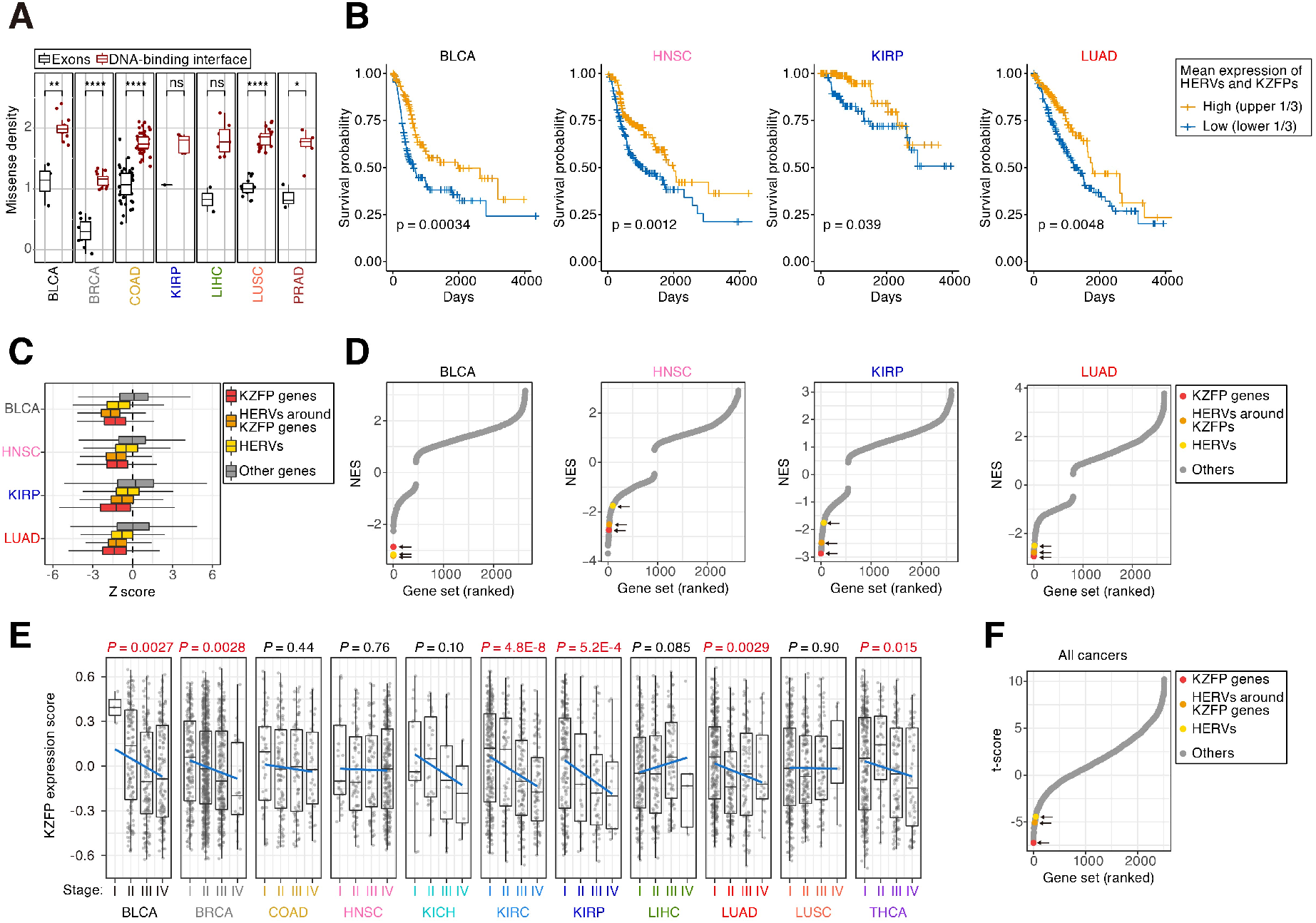
Association of the expression status of KZFPs and HERVs in tumors with cancer prognosis and progression. A) Accumulation of somatic missense mutations in the DNA-binding amino acid residues of KZFP genes. The mutation density (counts per mb per patient) of KZFP genes was compared between the DNA-binding amino acid residues (red) and the whole exonic regions (black). Results for KZFP genes with ≥1 mutations are shown. *P* values were calculated by two-sided Wilcoxon rank sum test. *, *P* < 0.05; **, *P* < 0.01; ****, *P* < 0.0001. B) Kaplan–Meier survival plots of cancer patients with high or low expression levels of HERVs and KZFPs. The results for BLCA, HNSC, KIRP, and LUAD tumors are shown (results for the other cancer types are shown in **Fig. S7A**). The stratification of the patients was according to the mean value of the gene set-wise expression scores (GSVA scores^46^) between KZFPs and HERVs. The results for the stratifications according to the GSVA scores of HERVs and KZFPs are shown in **Figs. S7B and S7C**, respectively. The *P* value was calculated by the two-sided log-rank test. C) Associations of respective genes and HERVs with the prognosis of cancer patients. The association was evaluated as the Z score in the Cox proportional hazards model, and the distributions of the Z score were compared among KZFPs, HERVs, HERVs around KZFPs (within 50 kb), and the other genes. Positive and negative Z scores indicate the associations with worse or better prognoses, respectively. D) Results of GSEA based on the Z scores in the Cox proportional hazards model. Positive and negative NES values indicate the associations with worse or better prognoses, respectively. Gene sets were ranked according to the NES value, and the gene sets of interest are highlighted. The high-scored gene sets are shown in **Fig. S8B**. E) Overall expression levels of KZFPs in respective cancer stages. The Y-axis indicates the GSVA score of KZFPs. The *P* value was calculated by single linear regression. F) Associations of the expression levels of respective gene sets with cancer progression. For each gene set, multiple linear regression analysis was performed with adjustment for cancer type-specific effects. Positive and negative t-scores indicate the tendencies of increase and decrease, respectively, in the GSVA scores along with cancer progression. Gene sets were ranked according to the t-score, and the gene sets of interest are highlighted. The high-scored gene sets are shown in **Fig. S9B**.

### Gene expression and phenotypic changes induced by the overexpression of KZFP genes in LUAD cells

The analysis of the chromatin immunoprecipitation-sequencing (ChIP-Seq) dataset of KZFPs (Imbeault et al.^33^) showed that many KZFPs preferentially bound to genes related to the cell cycle and cancer-associated signaling pathways, such as TGF-related pathways (TGF-β, BMP, SMAD2/3 pathways) and Wnt pathway (**Fig. S10**). These pathways are critical for the regulation of cell-matrix adhesion and are associated with cell migration/invasion and proliferation in cancers^34,35^. Notably, the expression levels of the genes related to the cell cycle and cell-matrix adhesion were negatively correlated with those of KZFP genes in tumors (**Fig. S3**) and associated with worse disease conditions (**Figs. S8B and S9B**), suggesting that KZFPs can modulate cancer phenotypes by altering the expression of these genes.

To assess the effects of elevated KZFP expression on cancer cells, we established a panel of A549 LUAD cells overexpressing 30 types of KZFPs (referred to as A549/KZFP cells) (**Fig. S11**) and subsequently investigated the phenotypic and gene expression changes caused by these KZFPs. Most of the tested KZFPs induced apoptosis (**Fig. 4A**), while many of the KZFPs suppressed cell growth, migration, and invasion (**Figs. 4B–D**). In total, the expression of 2,368 genes was altered by the overexpression of any of the tested KZFPs (**Fig. 4E**). Of note, the genes related to the cell cycle and cell-matrix adhesion were significantly downregulated by the overexpression of many types of KZFPs (**Fig. S12A**). Although the phenotypic and gene expression alterations caused by KZFPs were relatively similar among all types of A549/KZFP cells (**Figs. 4E and S12B**), these alterations were clearly associated (**Figs. S12C and S12D**), suggesting that the phenotypic changes in A549/KZFP cells were caused by alterations in gene expression. Overall, we demonstrated that a substantial fraction of KZFPs could suppress the phenotypes associated with cancer progression by altering gene expression in LUAD cells.

**Fig. 4.**
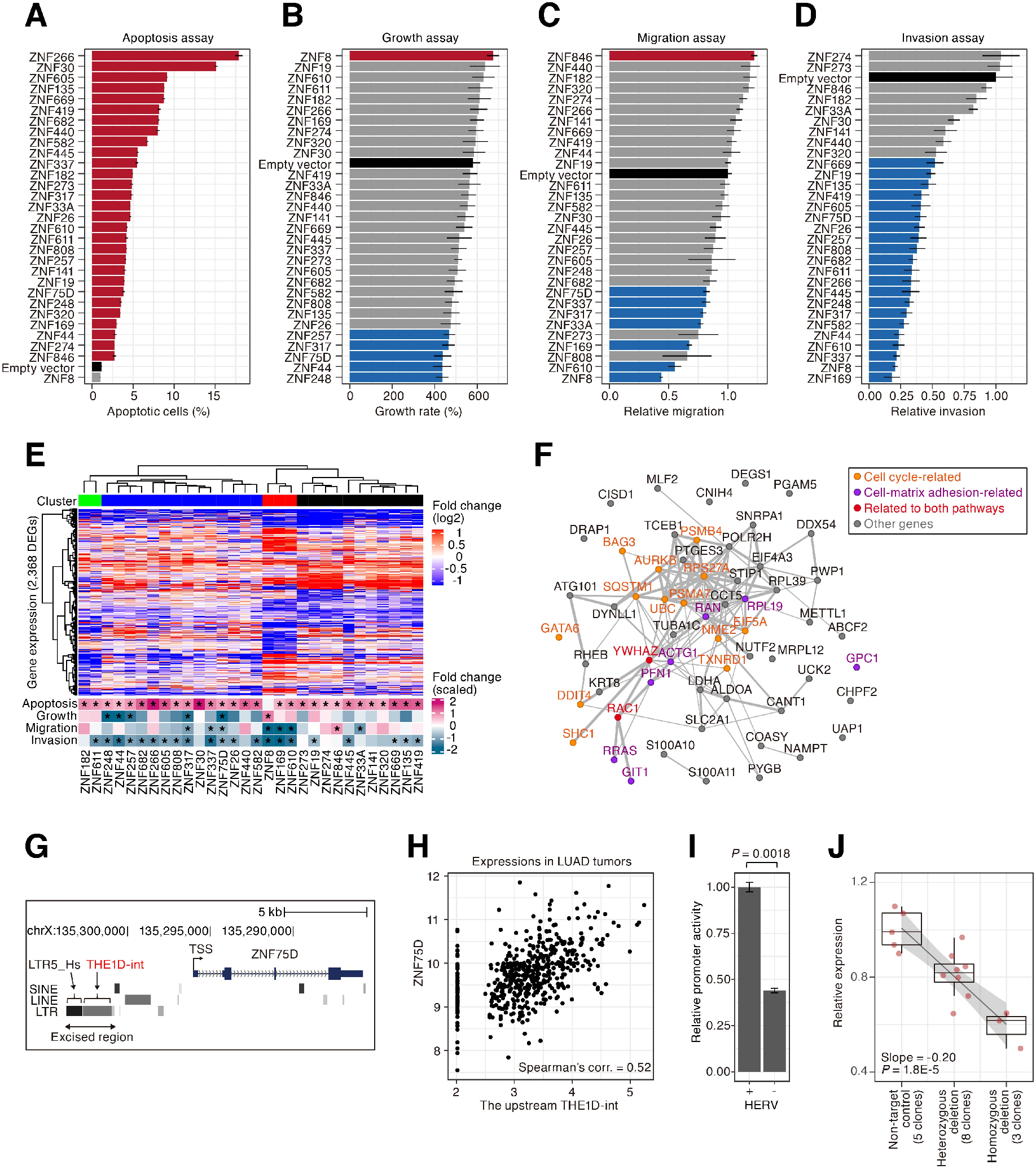
Phenotypic and gene expression changes caused by the overexpression of KZFPs in lung adenocarcinoma cells. A–D) Examinations of phenotypic changes in a panel of lung adenocarcinoma (A549) cells overexpressing 30 types of KZFPs (referred to as A549/KZFP cells). These 30 KZFPs satisfy the following criteria: 1) showing a positive correlation with the total expression of HERVs in tumors; 2) possessing expressed HERVs in the vicinity of its TSSs in tumors; and 3) having available ChIP-Seq data presented by a previous study (Imbeault et al.^33^). The results for the apoptosis assay (A), growth assay (B), migration assay (C), and invasion assay (D) are shown. The black bar indicates the result of the empty vector-transduced cells. Red or blue bars indicate the result of the cells in which the value significantly increased or decreased, respectively, compared to that in the empty vector-transduced cells in two-sided Student’s t-test (*P* value < 0.05). The error bar indicates the standard error of the mean (SEM). E) Phenotypic and gene expression changes in A549/KZFP cells. Upper) Heatmap showing the gene expression alterations of 2,368 differentially expressed genes (DEGs) identified in any of A549/KZFP cells compared to the empty vector-transduced cells. Gene expression-based clusters are indicated at the top of the heatmap. Lower) Heatmap summarizing the results of the experiments shown in A–D). For visualization, the values were log2-transformed and subsequently scaled (i.e., the standard deviation was adjusted at 1). An asterisk denotes a significant change in the value. F) Possible target genes of KZFPs critical for cancer progression. The details are described in **Fig. S13**. Edge indicates protein–protein interactions defined by the Search Tool for the Retrieval of Interacting Genes/Proteins (STRING) (version 11.0)^55^. The edge width represents the reliability score of the interaction. G) Schematic view of the *ZNF75D* gene locus. The region excised by CRISPR-Cas9 is indicated by the arrow. H) Expressional correlation between *ZNF75D* and the upstream THE1D-int in LUAD tumors. I) Effect of the HERV integrants on the promoter activity of *ZNF75D*. The effect was assessed by a luciferase reporter assay in A549 cells. A pair of the reporter plasmids harboring the *ZNF75D* promoters with and without these HERVs were constructed, and subsequently, the promoter activities were compared. Error bars indicate the SEM. *P* values were calculated by two-sided Student’s t-test. J) Effect of the CRISPR-Cas9 excision of these HERVs on the expression of *ZNF75D* in A549 cells. The mRNA expression level of *ZNF75D* in each clone of cells was measured by qRT-PCR. *P* values were calculated using linear regression.

To identify the target genes of KZFPs that are likely to be critical for cancer progression, we developed a scoring system for genes according to their expressional negative correlation with KZFPs, the association of their expression with worse clinical conditions, and their expressional suppression in A549/KZFP cells as well as considering the frequency of KZFP binding (**Fig. S13**). In this system, the high-scored genes included a substantial number of genes related to the cell cycle and cell-matrix adhesion (**Figs. 4F** and **S13D**). In particular, many genes related to cytoskeletal regulation (i.e., *ACTG1*, *GIT1*, *PFN1*, *RAC1*, and *RRAS*) that are critical for cell-matrix adhesion and modulate cell migration/invasion and proliferation^36^ were identified as targets of KZFPs. Additionally, a serine-threonine kinase gene (*AURKB*) and ubiquitin-proteasome pathway genes (*UBC*, *RPS27A*, *PSMB4*, and *PSMA7*) that are critical for cell cycle regulation^37,38^ were identified.

### Transcriptional modulation of cancer phenotype-associated KZFP genes by the adjacent HERVs in LUAD cells

ZNF75D was capable of altering all four investigated cancer phenotypes (**Figs. 4A–E**). In the region approximately 5 kb upstream of a TSS of *ZNF75D*, two HERV integrants (LTR5_Hs and THE1D-int) were present (**Fig. 4G**). THE1D-int was co-expressed with *ZNF75D* in LUAD tumors (**Fig. 4H**). A luciferase reporter assay showed that these two HERV elements exhibit enhancer activity in A549 cells (**Figs. 4I and S14**) regardless of their orientation (**Fig. S14E**). To test the significance of these HERVs on the transcriptional modulation of *ZNF75D*, we excised these two HERVs using the CRISPR-Cas9 system in A549 cells (**Fig. S6**) and demonstrated that the deletion of these HERVs decreased *ZNF75D* expression in an allelic number-dependent manner (**Fig. 4J**). These results suggest that these HERVs are involved in the transcriptional modulation of the *ZNF75D* in LUAD cells. Moreover, for 12 out of the 30 KZFP genes tested above, we investigated the transcriptional modulation potential of the adjacent HERVs by performing a luciferase reporter assay. HERVs in the vicinity of 7 KZFP genes (*ZNF141*, *ZNF248*, *ZNF30*, *ZNF320*, *ZNF44*, *ZNF611*, and *ZNF846*) enhanced the promoter activities of these genes in A549 cells (**Figs. S14F and S14G**). These results support the significance of HERVs in the transcriptional regulation of these KZFP genes in cancer cells.

## Discussion

In the present study, we found that the global activation of HERVs occurred in a substantial fraction of tumors (**Figs. 1E and S1C**). Although the ultimate cause of HERV activation in tumors is still unclear, the attenuation of the epigenetic silencing of HERVs would be related to HERV activation (**Figs. S1F and S1G**). HERV activation was associated with the synchronized induction of KZFP gene expression (**Figs. 1F and S2C**). Further analyses including *in vitro* experiments suggest that KZFPs are transcriptionally regulated by the adjacent HERVs (**Figs. 2 and 4G–4J**). Notably, the coordinated induction of KZFP expression was clearly associated with better disease conditions in multiple types of cancers (**Figs. 3B–F**). A substantial fraction of KZFPs could suppress the phenotypes related to tumor progression by altering gene expression in cultured cells (**Figs. 4A–4E**). These findings suggest that a repertoire of KZFPs cooperatively exerts suppressive effects on tumor progression. Collectively, we highlight the presence of tumor heterogeneity driven by the gene regulatory network comprising HERVs and KZFPs — the activation of HERVs in tumors induces the expression of adjacent KZFP genes, leading to the suppression of the progressive characteristics of cancers by altering gene expression.

Although our data highlight the significance of HERVs in the transcriptional regulation of KZFPs (**Fig. 2**), it is widely considered that one of the primary functions of KZFPs is the silencing of the disordered expression of TEs, including HERVs^28^. Such seemingly paradoxical findings suggest the presence of a transcriptional negative feedback loop between HERVs and KZFPs — once HERVs are derepressed globally, the regulatory activities of HERVs around KZFP genes are reactivated simultaneously, resulting in the induction of KZFP expression. In other words, KZFP genes seem to utilize HERVs as their regulatory sequences to detect the global derepression of HERVs. A previous report proposed the possibility that such negative feedback functions when the embryonic genome activation occurs to silence the activation of TEs including HERVs effectively^39^. This feedback system, at least regarding the induction of KZFP expression by HERVs, seems to work also in cancer cells and cause aberrant gene expression in tumors.

## Materials and Methods

### Ethical approval

The utilization of the TCGA multi-omics dataset was authorized by the National Cancer Institute (NCI) data access committee through the Database of Genotypes and Phenotypes (dbGaP; http://dbgap.ncbi.nlm.nih.gov) for the following projects: “Systematic identification of reactivated human endogenous retroviruses in cancers (#15126)”, “Effects of the genome-wide activation of human endogenous retroviruses on gene expression and cancer phenotypes (#18470)”, and “Screening of subclinical viral infections in healthy human tissues (#19481)”.

### Construction of the gene-HERV transcript model for RNA-Seq analysis

For the gene transcript model, GENCODE version 22 (for GRCh38/hg38) obtained from the GENCODE website (http://www.gencodegenes.org/) was used. For the HERV transcript model, the RepeatMasker output file (15-Jan-2014; for GRCh38/hg38) obtained from the UCSC genome browser (http://genome.ucsc.edu/) was used. From the gene model, transcripts with the flag “retained intron” were excluded. From the HERV model, HERV loci with low reliability scores (i.e., Smith-Waterman score < 2,500) were excluded. Additionally, the regions of HERV loci overlapping with the gene transcripts were also excluded. A gene-HERV transcript model was generated by concatenating the gene and HERV models. This model includes 60,483 protein-coding/non-coding genes in addition to 138,124 HERV loci, which occupy 3.4% of the genome.

### RNA-Seq data analysis of the TCGA dataset

Poly A-enriched RNA-Seq (mRNA-Seq) data provided by TCGA were analyzed. Of the RNA-Seq data, we analyzed only the data produced by pair-ended sequencing with a read length of 48–50 bp. The BAM-formatted read alignment file (for GRCh38/hg38) of the RNA-Seq data was downloaded from the Genomic Data Commons (GDC) data portal site (http://portal.gdc.cancer.gov/) using the GDC Data Transfer Tool (http://gdc.cancer.gov/access-data/gdc-data-transfer-tool/). To measure expression levels of HERVs and genes, RNA-Seq fragments mapped on HERVs and the exons of genes were counted using Subread featureCounts^40^ with the BAM file and the gene-HERV transcript model. The option “fracOverlap” was set at 0.25. The RNA-Seq fragments assigned to multiple features were not counted.

To control the quality of the RNA-Seq data used in the present study, we checked the proportion of non-assigned RNA-Seq fragments (i.e., the fragments that were uniquely mapped on the reference genome but not on HERVs or exons of genes) in each sequence library. For this proportion, outlier libraries were detected recursively using the Smirnov-Grubbs test (the threshold was set at 0.05). These outlier libraries were excluded from the downstream analyses. The final RNA-Seq data used in this study are summarized in **Data S4**.

The expression count matrices of the RNA-Seq data were separately prepared for the datasets of the respective types of cancers. In addition, the expression matrix including all tumor data was also prepared. Furthermore, the expression matrix, including the data from the tumors and corresponding normal adjacent tissues, was also prepared for each type of cancer. Genes and HERVs with low expression levels were removed from the expression matrix as follows. The counts per million (CPM) value of each gene and HERV locus were calculated in the respective RNA-Seq libraries. Subsequently, genes and HERVs were discarded from the expression matrix if the 90th percentile of CPM values was less than 0.2.

In each type of cancer, the expressed HERVs in tumors, which are HERVs included in the expression matrix of the corresponding types of cancers, were determined.

The total expression level of the HERVs was normalized as CPM. The expression levels of genes and HERV loci were normalized using variance-stabilizing transformation (VST) implemented in DESeq2 (version 1.18.1)^41^. This VST-normalized expression level was used unless otherwise noted.

### RNA-Seq data analysis of the CCLE dataset

The BAM-formatted read alignment file (for GRCh37/hg19) of the mRNA-Seq data was downloaded from the GDC data portal site (http://portal.gdc.cancer.gov/) using the GDC Data Transfer Tool (http://gdc.cancer.gov/access-data/gdc-data-transfer-tool/). The RNA-Seq data of CCLE used in this study are summarized in **Data S5**. Since the gene-HERV transcript model prepared above is for GRCh38/hg38, the genomic coordinates of the gene-HERV transcript model were converted to those in GRCh37/hg19 using UCSC liftOver (http://hgdownload.soe.ucsc.edu/admin/exe/linux.x86_64/liftOver). The option “minMatch” was set at 0.95. The generation of the expression count matrix, filtering of genes and HERVs with low expression levels, and normalization of the expression data were performed using the same procedures as those in the above section (“**RNA-Seq data analysis of the TCGA dataset**”).

### RNA-Seq analysis of A549/KZFP cells

The RNA-Seq sample information is summarized in **Data S6**. Low quality sequences in RNA-Seq fragments were trimmed using Trimmomatic (version 0.36)^42^ with the option “SLIDINGWINDOW:4:20”. RNA-Seq fragments were mapped to the human reference genome (GRCh38/hg38) using STAR (ver. 2.5.3a)^43^ with the gene-HERV transcript model. STAR was run using the same options and parameters as those used in the GDC mRNA Analysis Pipeline (https://docs.gdc.cancer.gov/Data/Bioinformatics_Pipelines/Expression_mRNA_Pipeline). The generation of the expression count matrix, filtering of genes and HERVs with low expression levels, and normalization of the expression data were performed using the same procedures as those in the above section (“**RNA-Seq data analysis of the TCGA dataset**”).

### Dimension reduction analysis of HERV expression profiles using t-distributed stochastic neighbor embedding (t-SNE)

The expression matrix including all tumor data was used in this analysis. The expression levels of the 1000 most highly expressed HERVs were used in the analysis. t-SNE analysis was performed using the “Rtsne” R package. For the analysis, the first 10 principle components of the HERV expression profiles were used, and the parameter “perplexity” was set at 70.

### ATAC-Seq data analysis

The ATAC-Seq data of tumors and normal adjacent tissues provided by TCGA (TCGA-ATAC_PanCan_Log2Norm_Counts.rds) was downloaded from the GDC website (https://gdc.cancer.gov/about-data/publications/ATACseq-AWG). This file contains the normalized read count matrix comprising all ATAC-Seq samples (n=796) and ATAC-Seq peaks (NFRs) (n=562,709) analyzed in the previous study^4^. In the respective types of cancers, the upper ¼ of NFRs with respect to the mean value were regarded as the NFRs that are active in the corresponding cancer types.

To calculate the fold enrichment of the overlaps between the expressed HERVs in tumors and NFRs, randomization-based enrichment analysis was performed as follows: genomic regions of NFRs were randomized using bedtools “shuffle”^44^ and subsequently, the number of NFRs on the expressed HERVs was counted. This process was repeated 1,000 times, and the mean value of the counts in the randomized datasets was regarded as the random expectation value. The fold enrichment was calculated by dividing the observed count by the random expectation value.

### DNA methylation data analysis

The DNA methylation data (produced by the methylation microarray HumanMethylation450 (Illumina)) of tumors and normal tissue controls were downloaded from the GDC data portal (http://portal.gdc.cancer.gov/) using the GDC Data Transfer Tool (http://gdc.cancer.gov/access-data/gdc-data-transfer-tool/). These data describe the methylation level (beta value; proportion of methylated CpGs at a CpG site) of each probe in the array. Probes overlapping with single nucleotide polymorphisms (SNPs) with >0.05 minor allele frequency were excluded from the analysis using the function “rmSNPandCH” implemented in the “DMRcate” library^45^ in R. The CpG sites that were on or proximal (<1 kb) to HERVs were extracted using the “slop” and “intersect” functions in bedtools^44^. DNA methylation data used in this study is summarized in **Data S7**.

### Preparation of gene sets for enrichment analyses

As sources of gene sets, “GO biological process”, “GO cellular component”, “MSigDB canonical pathway”, and “InterPro” were used. The gene sets in these sources were concatenated and used. “InterPro” is the collection of gene sets according to protein families or domains and includes the gene set “KRAB”, representing the KZFP family genes. “GO biological process” and “GO cellular component” were obtained from Gene Ontology (GO) consortium (http://geneontology.org/; GO validation date: 08/30/2017); “canonical pathway” was from MSigDB (http://software.broadinstitute.org/gsea/msigdb; version 6.1); and “InterPro” was from BioMart on the Ensembl website (https://www.ensembl.org; on 2/13/2018).

In addition, we defined the gene sets “HERVs” and “HERVs around KZFP genes”. The “HERV” gene set includes all expressed HERVs in tumors, while “HERVs around KZFP genes” includes the HERVs present in the genomic regions within 50 kb from the TSSs of KZFP genes expressed in tumors. These gene sets were used in **Figs. 3D, 3F, S8B, and S9B** in addition to the pre-defined gene sets.

Furthermore, we defined gene sets according to the expressional negative correlation with HERVs or KZFP genes as follows. In the respective tumor datasets of TCGA, Spearman’s correlations between the expression levels of respective genes and the total expression level of HERVs were calculated, and genes were ranked according to their median value in the datasets. The top 100, 200, and 500 genes with respect to their negative expressional correlation with HERVs were used as gene sets. Using the same procedures as above, the top 100, 200, and 500 genes with respect to the negative expressional correlation with KZFP genes were extracted and used as gene sets. As the representative value of KZFP expression, the gene set-wise expression score (Gene Set Variation Analysis (GSVA) score^46^) of the KZFP genes was used. The GSVA score is described in the following section (“**Calculation of the gene set-wise expression score using GSVA**”). These gene sets were used in **Fig. S10** in addition to the pre-defined gene sets.

### Calculation of the gene set-wise expression score using GSVA

The VST-normalized expression matrix was converted to the gene set-wise expression score matrix using GSVA^46^ with the gene sets prepared above. The option “minimum size of gene set” was set at 20.

### GSEA

To perform GSEA^29^, the R package “fgsea”^47^, a fast implementation of GSEA, was used. The parameters of “number of permutations” and “minimum size of gene set” were set at 10,000 and 50, respectively. In the analyses of **Figs. 1F and S4**, the Spearman’s correlations between the expression levels of respective genes and the total expression level of HERVs were used as statistical scores. In the analysis of **Fig. S3**, the Spearman’s correlations between the expression levels of respective genes and the GSVA score of the KZFP genes were used. In the analyses of **Figs. 3D and S8B**, Z scores in Cox proportional hazards regression were used (the Z score is described in the “**Survival analysis of the cancer patients**” section). In the analysis of **Fig. S12A**, Wald statistics of the respective genes in the differential expression analysis were used (the Wald statistic is described in the “**Differential expression analysis**” section).

### Summarizing the results of GSEA and GO enrichment analysis by removing redundant gene sets

Since the gene members of some gene sets highly overlapped with each other, redundant gene sets were removed from the results of the enrichment analyses as follows. Gene sets were ranked according to the score of interest (e.g., the mean value of normalized enrichment score (NES)). If the gene members of a certain gene set were highly overlapped with those of the upper-ranked gene sets, the gene set was removed from the result. As a statistic of the overlap, the Szymkiewicz–Simpson coefficient was used, and two gene sets were regarded as highly overlapped if the coefficient was greater than 0.7. This gene set filtering was applied to the analyses shown in **Figs. 1F, S8B, S9B, S10, and S12A**, which show only the top-ranked gene sets.

### GO enrichment analysis to identify gene sets that are preferentially present in the vicinity of the expressed HERVs

Randomization-based GO enrichment analysis was performed as follows. Only genes whose expression levels were detected in the TCGA tumor datasets were used. Regions of interest were defined as the regions within 50 kb from the TSSs of the gene members of a certain gene set. The genomic regions of HERVs were randomized using the “shuffle” function of bedtools^44^, and subsequently, the number of HERVs in the region of interest was counted. This process was repeated 1,000 times, and the mean value of the counts in the randomized datasets was regarded as the random expectation value. The fold enrichment was calculated by dividing the observed count by the random expectation value.

Additionally, we calculated the fold enrichments of HERVs in the regions within 10, 100, and 500 kb and 1 mb from the TSSs of the KZFP genes using the same procedures as above.

### Prediction of genes regulated by HERVs

The regulatory interactions between HERV loci and genes were predicted according to the following information: co-expression between HERVs and genes, positive correlations between HERV NFR activities and gene expression, negative correlations between HERV DNA methylation and gene expression, and pre-defined links between the regulatory sequences on HERVs and genes. The co-expression interaction was used in only pairs of HERVs and genes within 50 kb of each other, while the NFR–expression, methylation–expression, and pre-defined interactions were used in only pairs of HERVs and genes within 500 kb of each other. A co-expression interaction was defined if the expression of the HERV and gene were positively correlated (Spearman’s correlation > 0.4) in any type of cancer in TCGA. A methylation–expression interaction was defined if the DNA methylation level of the CpG site that is on or proximal (<1 kb) to a HERV and the expression of the gene were negatively correlated (Spearman’s correlation < −0.3) in any type of cancer or in the pan-cancer dataset in TCGA. As the source of NFR–expression interactions, the interactions defined in a previous study^4^ were used. As the source of pre-defined regulatory interactions, the interactions recorded in GeneHancer version 4.7 obtained from GeneLoc database (https://genecards.weizmann.ac.il/geneloc/index.shtml) were used.

### Mutation analysis

To define DNA-binding amino acids of KZFP genes, we first determined the precise genomic positions of KRAB and C2H2 zinc-finger domains as follows. For both of KRAB and C2H2 zinc-finger domains, Hidden Markov Model (HMM) profiles were generated using hmmbuild from HMMER2 [http://hmmer.org/]. Multiple sequence alignments used to build the HMM profiles were generated from the seed sequences downloaded from Pfam [https://academic.oup.com/nar/article/44/D1/D279/2503120]. Next, the human reference genome (GRCh37/hg19) was scanned using hmmpfam from HMMER2 with the built HMM profiles. The both strands of chromosomes translated in 3 reading frames were scanned. KZFP genes were collected if a KRAB domain had ≥2 downstream C2H2 zinc-fingers found on the same strand within 40kb, which corresponds to the maximum length from the first base of KRAB domain to the last base of zinc-finger domain. Detected KZFP genes were then annotated according to the Ensembl annotation (version 92; for GRCh37/hg19). Finally, the DNA-binding amino acid positions were inferred from C2H2 zinc-fingers annotated above, taking position 4th, 6th, 7th, and 10th (also called −1, +2, +3, and +6 positions) after the second cysteine of the C2H2. Only zinc-finger with a canonical C2H2 structure and associated with a KRAB domain was taken into account.

Processed mutation data were obtained from International Cancer Genome Consortium (ICGC) (release 27) (https://icgc.org/). Then we measured the somatic missense mutation density (counts per mb per patient) of KZFP genes in the DNA-binding amino acids and the whole exonic regions of the canonical transcript.

### Survival analysis of the cancer patients

The overall survival rate of the cancer patients was used for survival analyses with the R package “survival”. The survival curve of the patients was estimated by the Kaplan–Meier method, and statistical significance was evaluated by the two-sided log-rank test. With respect to the expression level of interest, the upper and lower third of patients were regarded as patients with higher and lower expression statuses, respectively. In **Figs. S7B and S7C**, the patients were stratified according to the GSVA expression scores of HERVs and KZFPs in tumors, respectively. In **Figs. 3B and S7A**, the patients were stratified according to the mean value of the GSVA scores of HERVs and KZFPs in tumors.

To examine the association of the expression level of each gene and HERV locus with the prognosis of cancer patients, Cox proportional hazards regression analysis was performed with adjustment for the effects of sex and race of the patients. In addition to HERVs, genes that were included in any of the gene sets prepared above were used.

### Association analysis of gene expression and cancer progression

Prostate adenocarcinoma (PRAD) tumors were excluded from the analysis since information on cancer stage for most PRAD patients was not available. In the analysis, cancer stage was regarded as an interval scale. For each type of cancer, the association between the expression of each gene and the progression of the cancer stage was evaluated by single linear regression. Similarly, the association between the GSVA score of each gene set and the progression of cancer stage for each type of cancer was evaluated using the same procedure. To evaluate the pan-cancer association of the GSVA score of each gene set and the progression of cancer stage, multiple linear regression analysis with adjustment for the effects of cancer type was performed.

### Analysis of a publicly available ChIP-Seq dataset of KZFPs

This analysis was based on a publicly available ChIP-Seq dataset of KZFPs in HEK293T cells presented in a previous study (Imbeault et al.^33^; GEO accession #: GSE78099). Information on pre-defied ChIP-Seq peaks (GSE78099_RAW.tar) was downloaded from the Gene Expression Omnibus (GEO) database (https://www.ncbi.nlm.nih.gov/geo/). Since these ChIP-Seq peaks (referred to as transcription factor binding sites; TFBSs) are for GRCh37/hg19, the genomic coordinates of these TFBSs were converted to those in GRCh38/hg38 using UCSC liftOver (http://hgdownload.soe.ucsc.edu/admin/exe/linux.x86_64/liftOver). The option “minMatch” was set at 0.95. If multiple technical replicates of ChIP-Seq are available for one KZFP, the replicate files were merged using the bedtools “merge”^44^ with the options “-c 5 -o mean”. KZFPs were removed from the downstream analyses if the total number of TFBSs was less than 500. If >10,000 TFBSs were available for one KZFP, only the high-scored 10,000 TFBSs were used for the analyses.

To identify sets of genes that are preferentially targeted by a certain KZFP, genomic region enrichment analysis (GREAT)^48^ was performed as follows. Only genes whose expression levels were detected in the TCGA tumor datasets were used. Regions of interest were defined as the regions within 10 kb from the TSSs of the gene members of a certain gene set. Regions of background were defined as the regions within 10 kb from the TSSs of genes belonging to any of the gene sets. The lengths of the regions of interest and regions of background were calculated and referred to as L_i_ and L_b_, respectively. In the regions of interest and regions of background, the numbers of TFBSs were counted (referred to as counts of interest (C_i_) and counts of background (C_b_), respectively). The fold enrichment value was calculated by dividing C_i_/C_b_ by L_i_/L_b_, and the statistical significance was evaluated using a binomial test.

### Differential expression analysis

Differential expression analysis was performed using DESeq2 (version 1.18.1)^41^ in R. Genes that were included in any of the gene sets prepared above were used. A549/KZFP cells and empty vector-transduced cells was compared (**Fig. 4E**). Additionally, comparison was conducted between A549 cells in which HERV-enhancer1 were excised versus the non-target control cells (**Fig. 2G**). Statistical significance was evaluated by the Wald test with false discovery rate (FDR) correction using the Benjamini-Hochberg (BH) method.

### Scoring system of genes for predicting the targets of KZFPs critical for cancer progression

The scheme is summarized in **Fig. S13A**. For each gene, the following scores were defined. The TCGA expressional correlation score was defined as the Spearman’s correlation between the expression of each gene and GSVA score of KZFPs in the TCGA dataset (the median value among all cancer types was used). The CCLE expressional correlation score was also defined using the same procedure but on the CCLE dataset. The prognosis score was defined as the Z score representing the association of each gene with the prognosis of cancer patients (the mean value among BLCA, HNSC, KIRP, and LUAD tumors was used). This Z score was described in the above section “**Survival analysis of the cancer patients**”. The progression score was defined as the t-score representing the association of each gene with cancer progression (the mean value among BLCA, BRCA, KIRC, KIRP, LUAD, and thyroid carcinoma (THCA) tumors was used). This t-score was described in the above section “**Association analysis of gene expression and cancer progression**”. The suppression score was defined as the mean value of the Wald scores in the differential expression analysis among the A549/KZFP cells. This Wald score was described in the above section “**RNA-Seq analysis of A549/KZFP cells**”. Regarding the TCGA and CCLE correlation scores and suppression scores, the signs of the scores were inverted. All scores were standardized as Z scores and subsequently quantile-normalized. Genes were extracted if the minimum score was greater than 0.5 and the median score was greater than 1. Of the extracted genes, genes targeted by ≥10 KZFPs were further extracted and regarded as the target genes of KZFPs critical for cancer progression. A gene was regarded as the target of a certain KZFP if the KZFP bound to the regions within 10 kb from the TSSs of the gene. In this analysis, only TSSs of “principal transcripts” (Principals 1–3) defined by APPRIS^49^ were used. If >1,000 genes were assigned to a certain KZFP as its targets, only the top 1,000 genes having high-scored TFBSs were used.

### Data visualization

All visualizations were performed in R. Graphs were plotted using the “ggplot2” package or the pre-implemented function “plot” unless otherwise noted. Heatmaps were drawn using the “ComplexHeatmap” package^50^. Networks were plotted using the “igraph” package. Kaplan–Meier plots were drawn using the “ggsurvplot” function in the “survminer” package.

### Cell culture

HEK293T cells (CRL-11268; ATCC, Manassas, VA) were cultured in Dulbecco’s modified Eagle’s medium (Sigma-Aldrich, St .Louis, MO; #D6046) with 10% fetal bovine serum (FBS; Sigma-Aldrich #172012-500ML) and 1% penicillin streptomycin (Sigma-Aldrich #P4333-100ML). A549 cells (CCL-185; ATCC) were cultured in Ham’s F-12K (Kaighn’s) medium (Thermo Fisher Scientific, Waltham, MA; #21127022) with 10% FBS (guaranteed doxycycline free; Thermo Fisher Scientific; #2023-03) and 1% penicillin streptomycin. A459/KZFP cells were cultured in F-12K medium with 1.0 μg/ml puromycin (Invivogen, San Diego, CA; #ant-pr-1). An A549 cell line stably expressing Cas9 (A549/Cas9 cells) was cultured in F-12K medium with 10% FBS (guaranteed doxycycline free; Thermo Fisher Scientific; #2023-03) and 5.0 μg/ml blasticidin (Invivogen #ant-bl-1). All cells were cultured in 5% CO_2_ at 37°C.

### Establishment of a panel of A549/KZFP cells

A549 cells were selected as the parental cells since the expression levels of KZFPs (and HERVs) were relatively low in this cell line (**Fig. S11A**). We selected 30 types of KZFP genes satisfying the following criteria: 1) showing a positive correlation (Spearman’s correlation > 0.3) between its expression and the total expression of HERVs in >2 types of cancers; 2) possessing expressed HERVs within the vicinity (<20 kb) of its TSSs in tumors; 3) showing a positive correlation (Spearman’s correlation > 0.3) between its expression and the expression of HERV loci in the vicinity (<20 kb) of its TSSs in >2 types of cancers; 4) having available ChIP-Seq data presented by a previous study (Imbeault et al.^33^). Information of the selected KZFP genes is summarized in **Data S8**.

To prepare the lentiviral vectors expressing x3 HA-tagged KZFPs, HEK293T cells were co-transfected with 12 μg of pCAG-HIVgp (RDB04394, kindly provided by Dr. Hiroyuki Miyoshi), 10 μg of pCMV-VSV-G-RSV-Rev (RDB04393, kindly provided by Dr. Hiroyuki Miyoshi), and 17 μg of pEXPpSIN-TRE-GW ZNF-3xHA^33^ by the calcium phosphate method. The pEXPpSIN-TRE-GW ZNF-3xHA plasmids encode respective HA-tagged KZFP proteins. After 12 hours of transfection, the culture medium was changed to fresh F-12K medium. After 48 hours of transfection, the culture supernatant including lentivector particles was collected. A549 cells were infected with these particles at a multiplicity of infection (MOI) of 0.1. After 2 days of infection, the cells were selected with puromycin (1 μg/ml) for 7 days. Three days before the start of the experiments, doxycycline (1.0 μg/ml) was added to induce the expression of KZFP. The expression of KZFP was verified by western blotting with the HA-specific antibody (Roche, Basel, Switzerland; #12013819001). Empty vector-transduced A549 cells (referred to as negative control cells (NC cells)) were established according to the procedures described above.

### Apoptosis detection assay

A549/KZFP cells and NC cells were stained with Annexin V conjugated to Alexa Fluor™ 647 (Invitrogen Carlsbad, CA; #S32357). After staining, the number of Annexin V-positive cells was counted by a FACSCalibur system (BD Biosciences, San Jose, CA), and the rate of apoptotic cells was calculated. A single set of triplicate experiments was performed, and the mean and standard error of the mean (SEM) values are shown in **Fig. 4A**. Statistical tests were performed by two-sided Student’s t-test with a threshold of 0.05.

### Cell growth assay

A549/KZFP cells and NC cells were seeded at 1.0 × 10^5^ cells/well in 6-well plates (Thermo Fisher Scientific). After 72 hours of seeding, the number of cells was counted manually under a microscope, and the growth rate of the cells was calculated. Single-replicate experiments were performed at least 7 times independently, and the mean and SEM values are shown in **Fig. 4B**. Statistical tests were performed by two-sided Student’s t-test with a threshold of 0.05.

### Cell scratch assay (wound-healing assay^51^)

A549/KZFP cells and NC cells were seeded in 12-well plates (Thermo Fisher Scientific) and cultured until >90% confluence. A single straight wound was formed in each well by scratching with a sterile 1,000 μl pipette tip. The cells were washed with phosphate-buffered saline (PBS), and 2 ml of F-12K medium was added. Images were taken under a microscope immediately after the scratch and again after 24 hours. Using ImageJ^52^ with in-house scripts, the area (pixels) in which cells migrated for 24 hours was calculated. Triplicate experiments were performed independently twice. Regarding the mean and SEM, the average values between the two sets of experiments are shown in **Fig. 4C**. Statistical tests were performed by two-sided Student’s t-test in each set of experiments with a threshold of 0.05. Only if a significant difference was observed in both sets of experiments, the comparison was considered significant.

### Cell invasion assay

An invasion assay was performed using a 96-well Transwell plate (8.0-μm pore size) (Corning, Corning, NY #3374) with Corning Matrigel Basement Membrane Matrix (Corning #354234). The Matrigel matrix was diluted 50-fold with serum-free F-12K medium. To coat the Transwell insert plate, 30 μl of Matrigel matrix was dispensed into the insert plate. After 2 hours of incubation, 20 μl of the supernatant was removed from the coated Transwell plate. Subsequently, A549/KZFP cells and NC cells were seeded at 5.0 × 10^4^ cells/well in the insert plate. The insert plate was filled with serum-free F-12K medium, while the reservoir plate was filled with F-12K medium with 10% FBS. After incubation at 37°C for 48 hours, the cells that had invaded the Matrigel and migrated to the opposite side of the insert plate were washed with PBS, stripped with Trypsin-EDTA, and stained with calcein AM (Invitrogen #C3100MP). To evaluate the degree of cell invasion, the fluorescence intensity of the cells was measured using a 2030 ARVO X multi-label counter (PerkinElmer, Waltham, MA). The relative fluorescence intensity was calculated as (FI_i_ - FI_b_) / (FI_c_ - FI_b_), where FI_i_ denotes the fluorescence intensity of the A549/KZFP cells of interest, FI_b_ denotes the intensity of the blank, and FI_c_ denotes the intensity of NC cells. Triplicate experiments were performed independently twice. Regarding the mean and SEM, the average values between the two sets of experiments are shown in **Fig. 4D**. Statistical tests were performed by two-sided Student’s t-test in each set of experiments with a threshold of 0.05. Only if a significant difference was observed in both sets of experiments, the comparison was considered significant.

### Construction of plasmids for the luciferase reporter assay

Genomic DNA from the human peripheral blood lymphocytes of a healthy donor was used as the DNA source. A luciferase reporter vector, pGL3-basic (Promega, Madison, WI), was used. Using nested PCR, the genomic region indicated by the arrow in **Figs. S14A–14B** was cloned into pGL3-basic.

Information on the plasmids and primers prepared in this section is summarized in **Data S9 and S10**, respectively.

### Luciferase reporter assay to assess the promoter activity of genes

A549 cells were seeded at 1.0 × 10^5^ cells/well in 12-well plates (Thermo Fisher Scientific). After 24 hours of seeding, the luciferase reporter plasmid was transfected using polyethylenimine transfection. To fairly compare the reporter activities of the two plasmids with different sequence lengths, 1 μg of the longer plasmid and the same molar of the shorter plasmid were used for the transfection. After 12 hours of transfection, the culture medium was changed to fresh F-12K medium. After 48 hours of transfection, the luminescence intensity of the transfected cells was measured using a 2030 ARVO X multi-label counter (PerkinElmer) or a GloMax® Explorer Multimode Microplate Reader 3500 (Promega) with a BrillianStar-LT assay system (Toyo-b-net, Tokyo, Japan; #307-15373 BLT100). A single set of triplicate experiments was performed, and the mean and SEM values are shown in **Figs. 4I and S14E–G**. Statistical tests were performed by two-sided Student’s t-test with a threshold of 0.05.

### Establishment of HERV-excised cells

First, an A549 cell line stably expressing Cas9 (referred to as A549/Cas9 cells) was established as follows. To prepare the lentiviral vectors expressing Cas9, HEK293T cells were co-transfected with 12 μg of pCAG-HIVgp, 10 μg of pCMV-VSV-G-RSV-Rev, and 17 μg of plentiCas9-Blast (Addgene, Watertown, MA; #52962) by the calcium phosphate method. After 12 hours of transfection, the culture medium was changed to fresh F-12K medium. After 48 hours of transfection, the culture supernatant including lentivector particles was collected. A549 cells were infected with these particles at an MOI of 0.1. After 2 days of infection, the cells were selected with blasticidin (5 μg/ml) for 7 days. After selection, single cell clones were obtained through the limiting dilution method. By screening the expression level of Cas9 among the candidate clones, A549/Cas9 cells were established.

To excise the target HERV, a pair of guide RNAs (gRNAs) were designed in the upstream and downstream regions of the HERV using the web applications of sgRNA designer^53^ (http://portals.broadinstitute.org/gpp/public/analysis-tools/sgrna-design) or CRISPOR^54^ (http://crispor.tefor.net). The gRNA information is summarized in **Data S11**. The gRNA was cloned into a gRNA expression plasmid, lentiGuide-Puro (Addgene #52963). A pair of gRNA expression plasmids was co-transfected into the A549/Cas9 cells by electroporation using the NEON Transfection System (ThermoFisher) (1200 V; 30 ms; 2 times pulse; 1.0 × 10^5^ cells; and 500 ng of each plasmid). After transfection, the cells were selected with 1 μg/ml puromycin for 3 days. After selection, single cell clones were obtained through the limiting dilution method. Of these candidate clones, the clones in which homozygous or heterozygous excision of the target HERV occurred were screened using PCR (**Fig. S6**).

Regarding homozygous clones, the PCR fragments were checked through molecular cloning into a TOPO vector (Invitrogen #450245) followed by Sanger’s sequencing.

### qRT-PCR

Total RNA was extracted from cells by the QIAamp RNA Blood Mini Kit (QIAGEN, Hilden, Germany; # 52304) and subsequently treated with DNase I, Amplification Grade (Invitrogen #18068015). cDNA was synthesized by reverse transcription of the total RNA using SuperScript III reverse transcriptase (Life technologies #18080044) with Oligo(dT)12-18 Primer (Invitrogen #18418012). qRT-PCR was performed on the cDNA using a CFX Connect Real-Time PCR Detection System (Bio-Rad, Richmond, CA; #1855201J1) with a TaqMan® Gene Expression Assay kit (Thermo Fisher Scientific). The primer and TaqMan probe information are listed in **Data S12**. *GAPDH* was used as an internal control.

### Preparation of RNA-Seq samples and sequencing

Cells were seeded at 1.0 × 10^6^ cells in 100 mm dishes (Thermo Fisher Scientific EasYDish #150466). After 48 hours of seeding, the cells were harvested and stored at −80°C. Total RNA was extracted from the cells by the QIAamp RNA Blood Mini Kit (QIAGEN #52304) and subsequently treated with RNase-Free DNase Set (QIAGEN #79254).

Quality checks, library construction, and sequencing were performed by Novogene (https://en.novogene.com). Pair-end 150-bp read length sequencing was performed on an Illumina NovaSeq 6000 system.

### Code availability

Computer codes used in the present study will be available in the GitHub repository (https://github.com/TheSatoLab/HERV_Pan-cancer_analysis).

### Data availability

RNA-seq data reported in this paper will be available in GEO (https://www.ncbi.nlm.nih.gov/geo/; GSE141803).

## Acknowledgments

We would like to thank Naoko Misawa (Institute for Frontier Life and Medical Sciences, Kyoto University, Japan) and Kyoichiro Nomura (Yamaguchi University, Japan) for technical supports; Julien Pontis (School of Life Sciences, Ecole Polytechnique Federale de Lausanne (EPFL), Switzerland), Shohei Kojima and Junna Kawasaki (Institute for Frontier Life and Medical Sciences, Kyoto University, Japan), and Shiro Yamada (Tokai University, Japan) for thoughtful comments. The super-computing resource, SHIROKANE, was provided by Human Genome Center, The Institute of Medical Science, The University of Tokyo, Japan. The results shown here are in part based upon data generated by the TCGA Research Network (https://www.cancer.gov/tcga). **Funding:**This study was supported in part by AMED J-PRIDE JP19fm0208006 (to K.S.); AMED Research Program JP19fk0410014 (to Y.K. and K.S.) and JP19fk0410019 (to K.S.); JST CREST (to K.S.); JSPS KAKENHI Scientific Research B JP18H02662 (to K.S.); JSPS KAKENHI Scientific Research on Innovative Areas JP16H06429 (to K.S.), JP16K21723 (to K.S.), JP17H05813 (to K.S.), and JP19H04826 (to K.S.); JSPS Research Fellow PD JP19J01713 (to J.I.) and DC1 JP19J20488 (to I.K.); JSPS Core-to-Core program (A. Advanced Research Networks) (to Y.K. and K.S.); Joint Usage/Research Center program of Institute for Frontier Life and Medical Sciences, Kyoto University (to K.S.); Takeda Science Foundation (to K.S.); ONO Medical Research Foundation (to K.S.); Ichiro Kanehara Foundation (to K.S.); Lotte Foundation (to K.S.); and Mochida Memorial Foundation for Medical and Pharmaceutical Research (to K.S.).

## Author contributions

J.I. conceived the study; J.I. and A.C. mainly performed bioinformatics analyses; I.K., H.N., I.I., P.T., and D.T. supported bioinformatics analyses; I.K mainly performed experimental analyses; A.S. and Y.K. supported experimental analyses; Y.K., P.T., and D.T. provided reagents; J.I., I.K., and K.S. prepared the figures; J.I., I.K., and K.S. wrote the initial draft of the manuscript; all authors contributed to data interpretation, designed the research, revised the paper, and approved the final manuscript.

## Declaration of Interests

The authors declare that they have no competing interests.

## Notes

**Conflict of interest:** The authors declare that no competing interests exist.

## References

1 Weinstein, J. N. et al. The Cancer Genome Atlas Pan-Cancer analysis project. Nat Genet 45, 1113–1120, doi:10.1038/ng.2764 (2013).

2 Uhlen, M. et al. A pathology atlas of the human cancer transcriptome. Science 357, doi:10.1126/science.aan2507 (2017).

3 Bradner, J. E., Hnisz, D. & Young, R. A. Transcriptional Addiction in Cancer. Cell 168, 629–643, doi:10.1016/j.cell.2016.12.013 (2017).

4 Corces, M. R. et al. The chromatin accessibility landscape of primary human cancers. Science 362, doi:10.1126/science.aav1898 (2018).

5 Chen, H. et al. A Pan-Cancer Analysis of Enhancer Expression in Nearly 9000 Patient Samples. Cell 173, 386–399.e312, doi:10.1016/j.cell.2018.03.027 (2018).

6 Chuong, E. B., Elde, N. C. & Feschotte, C. Regulatory activities of transposable elements: from conflicts to benefits. Nat Rev Genet 18, 71–86, doi:10.1038/nrg.2016.139 (2017).

7 Coffin, J. M. Retroviruses. (Cold Spring Harbor Laboratory Press, 2002).

8 Lander, E. S. et al. Initial sequencing and analysis of the human genome. Nature 409, 860–921, doi:10.1038/35057062 (2001).

9 Sundaram, V. et al. Widespread contribution of transposable elements to the innovation of gene regulatory networks. Genome Res 24, 1963–1976, doi:10.1101/gr.168872.113 (2014).

10 Ito, J. et al. Systematic identification and characterization of regulatory elements derived from human endogenous retroviruses. PLoS Genet 14, e1006883, doi:10.1371/journal.pgen.1006883 (2017).

11 Kunarso, G. et al. Transposable elements have rewired the core regulatory network of human embryonic stem cells. Nat Genet 42, 631–634, doi:10.1038/ng.600 (2010).

12 Pi, W. et al. Long-range function of an intergenic retrotransposon. Proc Natl Acad Sci U S A 107, 12992–12997, doi:10.1073/pnas.1004139107 (2010).

13 Emera, D. et al. Convergent evolution of endometrial prolactin expression in primates, mice, and elephants through the independent recruitment of transposable elements. Mol Biol Evol 29, 239–247, doi:10.1093/molbev/msr189 (2012).

14 Wang, J. et al. Primate-specific endogenous retrovirus-driven transcription defines naive-like stem cells. Nature 516, 405–409, doi:10.1038/nature13804 (2014).

15 Ferreira, L. M. et al. A distant trophoblast-specific enhancer controls HLA-G expression at the maternal-fetal interface. Proc Natl Acad Sci U S A 113, 5364–5369, doi:10.1073/pnas.1602886113 (2016).

16 Zhang, Y. et al. Transcriptionally active HERV-H retrotransposons demarcate topologically associating domains in human pluripotent stem cells. Nat Genet 51, 1380–1388, doi:10.1038/s41588-019-0479-7 (2019).

17 Chuong, E. B., Elde, N. C. & Feschotte, C. Regulatory evolution of innate immunity through co-option of endogenous retroviruses. Science 351, 1083–1087, doi:10.1126/science.aad5497 (2016).

18 Slotkin, R. K. & Martienssen, R. Transposable elements and the epigenetic regulation of the genome. Nat Rev Genet 8, 272–285, doi:10.1038/nrg2072 (2007).

19 Deniz, O., Frost, J. M. & Branco, M. R. Regulation of transposable elements by DNA modifications. Nat Rev Genet 20, 417–431, doi:10.1038/s41576-019-0106-6 (2019).

20 Rooney, M. S., Shukla, S. A., Wu, C. J., Getz, G. & Hacohen, N. Molecular and genetic properties of tumors associated with local immune cytolytic activity. Cell 160, 48–61, doi:10.1016/j.cell.2014.12.033 (2015).

21 Smith, C. C. et al. Endogenous retroviral signatures predict immunotherapy response in clear cell renal cell carcinoma. J Clin Invest 128, 4804–4820, doi:10.1172/jci121476 (2018).

22 Solovyov, A. et al. Global Cancer Transcriptome Quantifies Repeat Element Polarization between Immunotherapy Responsive and T Cell Suppressive Classes. Cell Rep 23, 512–521, doi:10.1016/j.celrep.2018.03.042 (2018).

23 Panda, A. et al. Endogenous retrovirus expression is associated with response to immune checkpoint blockade in clear cell renal cell carcinoma. JCI Insight 3, doi:10.1172/jci.insight.121522 (2018).

24 Attig, J. et al. LTR retroelement expansion of the human cancer transcriptome and immunopeptidome revealed by de novo transcript assembly. Genome Res 29, 1578–1590, doi:10.1101/gr.248922.119 (2019).

25 Babaian, A. & Mager, D. L. Endogenous retroviral promoter exaptation in human cancer. Mob DNA 7, 24, doi:10.1186/s13100-016-0080-x (2016).

26 Jang, H. S. et al. Transposable elements drive widespread expression of oncogenes in human cancers. Nat Genet 51, 611–617, doi:10.1038/s41588-019-0373-3 (2019).

27 Hutter, C. & Zenklusen, J. C. The Cancer Genome Atlas: Creating Lasting Value beyond Its Data. Cell 173, 283–285, doi:10.1016/j.cell.2018.03.042 (2018).

28 Ecco, G., Imbeault, M. & Trono, D. KRAB zinc finger proteins. Development 144, 2719–2729, doi:10.1242/dev.132605 (2017).

29 Subramanian, A. et al. Gene set enrichment analysis: a knowledge-based approach for interpreting genome-wide expression profiles. Proc Natl Acad Sci U S A 102, 15545–15550, doi:10.1073/pnas.0506580102 (2005).

30 Ghandi, M. et al. Next-generation characterization of the Cancer Cell Line Encyclopedia. Nature 569, 503–508, doi:10.1038/s41586-019-1186-3 (2019).

31 Huntley, S. et al. A comprehensive catalog of human KRAB-associated zinc finger genes: insights into the evolutionary history of a large family of transcriptional repressors. Genome Res 16, 669–677, doi:10.1101/gr.4842106 (2006).

32 Fishilevich, S. et al. GeneHancer: genome-wide integration of enhancers and target genes in GeneCards. Database (Oxford) 2017, doi:10.1093/database/bax028 (2017).

33 Imbeault, M., Helleboid, P. Y. & Trono, D. KRAB zinc-finger proteins contribute to the evolution of gene regulatory networks. Nature 543, 550–554, doi:10.1038/nature21683 (2017).

34 Horbelt, D., Denkis, A. & Knaus, P. A portrait of Transforming Growth Factor beta superfamily signalling: Background matters. Int J Biochem Cell Biol 44, 469–474, doi:10.1016/j.biocel.2011.12.013 (2012).

35 Wang, Y. Wnt/Planar cell polarity signaling: a new paradigm for cancer therapy. Mol Cancer Ther 8, 2103–2109, doi:10.1158/1535-7163.Mct-09-0282 (2009).

36 Hall, A. The cytoskeleton and cancer. Cancer Metastasis Rev 28, 5–14, doi:10.1007/s10555-008-9166-3 (2009).

37 Dar, A. A., Goff, L. W., Majid, S., Berlin, J. & El-Rifai, W. Aurora kinase inhibitors--rising stars in cancer therapeutics? Mol Cancer Ther 9, 268–278, doi:10.1158/1535-7163.Mct-09-0765 (2010).

38 Nakayama, K. I. & Nakayama, K. Ubiquitin ligases: cell-cycle control and cancer. Nat Rev Cancer 6, 369–381, doi:10.1038/nrc1881 (2006).

39 Pontis, J. et al. Hominoid-Specific Transposable Elements and KZFPs Facilitate Human Embryonic Genome Activation and Control Transcription in Naive Human ESCs. Cell Stem Cell 24, 724–735.e725, doi:10.1016/j.stem.2019.03.012 (2019).

40 Liao, Y., Smyth, G. K. & Shi, W. featureCounts: an efficient general purpose program for assigning sequence reads to genomic features. Bioinformatics 30, 923–930, doi:10.1093/bioinformatics/btt656 (2014).

41 Love, M. I., Huber, W. & Anders, S. Moderated estimation of fold change and dispersion for RNA-seq data with DESeq2. Genome Biol 15, 550, doi:10.1186/s13059-014-0550-8 (2014).

42 Bolger, A. M., Lohse, M. & Usadel, B. Trimmomatic: a flexible trimmer for Illumina sequence data. Bioinformatics 30, 2114–2120, doi:10.1093/bioinformatics/btu170 (2014).

43 Dobin, A. et al. STAR: ultrafast universal RNA-seq aligner. Bioinformatics 29, 15–21, doi:10.1093/bioinformatics/bts635 (2013).

44 Quinlan, A. R. & Hall, I. M. BEDTools: a flexible suite of utilities for comparing genomic features. Bioinformatics 26, 841–842, doi:10.1093/bioinformatics/btq033 (2010).

45 Peters, T. J. et al. De novo identification of differentially methylated regions in the human genome. Epigenetics Chromatin 8, 6, doi:10.1186/1756-8935-8-6 (2015).

46 Hanzelmann, S., Castelo, R. & Guinney, J. GSVA: gene set variation analysis for microarray and RNA-seq data. BMC Bioinformatics 14, 7, doi:10.1186/1471-2105-14-7 (2013).

47 Sergushichev, A. A. Fast gene set enrichment analysis. bioRxiv, doi:10.1101/060012 (2016).

48 McLean, C. Y. et al. GREAT improves functional interpretation of cis-regulatory regions. Nat Biotechnol 28, 495–501, doi:10.1038/nbt.1630 (2010).

49 Rodriguez, J. M. et al. APPRIS: annotation of principal and alternative splice isoforms. Nucleic Acids Res 41, D110–117, doi:10.1093/nar/gks1058 (2013).

50 Gu, Z., Eils, R. & Schlesner, M. Complex heatmaps reveal patterns and correlations in multidimensional genomic data. Bioinformatics 32, 2847–2849, doi:10.1093/bioinformatics/btw313 (2016).

51 Liang, C. C., Park, A. Y. & Guan, J. L. In vitro scratch assay: a convenient and inexpensive method for analysis of cell migration in vitro. Nat Protoc 2, 329–333, doi:10.1038/nprot.2007.30 (2007).

52 Schneider, C. A., Rasband, W. S. & Eliceiri, K. W. NIH Image to ImageJ: 25 years of image analysis. Nat Methods 9, 671–675 (2012).

53 Sanson, K. R. et al. Optimized libraries for CRISPR-Cas9 genetic screens with multiple modalities. Nat Commun 9, 5416, doi:10.1038/s41467-018-07901-8 (2018).

54 Concordet, J. P. & Haeussler, M. CRISPOR: intuitive guide selection for CRISPR/Cas9 genome editing experiments and screens. Nucleic Acids Res 46, W242–w245, doi:10.1093/nar/gky354 (2018).

55 Szklarczyk, D. et al. STRING v11: protein-protein association networks with increased coverage, supporting functional discovery in genome-wide experimental datasets. Nucleic Acids Res 47, D607–d613, doi:10.1093/nar/gky1131 (2019).

