## Supplementary figures for "Endogenous retroviruses drive KRAB zinc-finger family protein expression for tumor suppression"

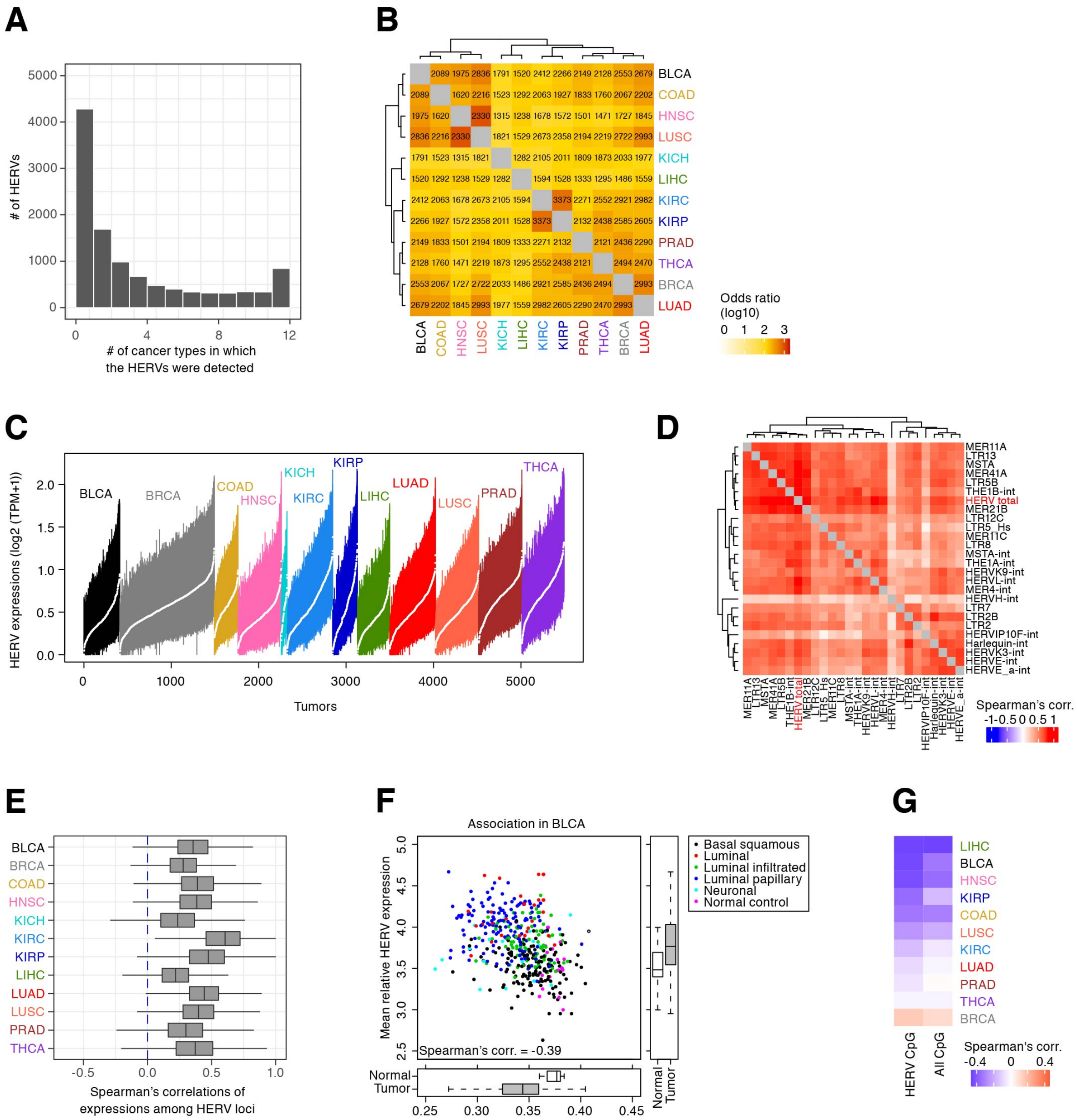


**Fig. S1 Characterization of HERV expression in 12 types of solid cancers.**

A) Histogram showing the numbers of cancer types in which respective expressed HERV loci were detected.

B) Pairwise commonalities of the sets of expressed HERVs among cancer types. In the heatmap, the color indicates the degree of overlap (log10 (odds ratio)), and the numerical characters indicate the number of expressed HERVs that were commonly detected in a pair of cancer types.

C) Boxplot showing the distribution of the expression levels of the respective HERV loci in each tumor. The Y-axis indicates the expression levels of the respective HERV loci (log2 (transcripts per million (TPM) + 1)). A colored line and a white dot indicate the inter-quantile range and the median value, respectively. Tumors were ordered according to the cancer type and the median value.

D) Pairwise expression correlations among HERV groups in BLCA tumors. The results for the 25 most highly expressed HERV groups are shown. In addition, the result of the total expression level of HERVs is also included.

E) Distribution of the pairwise expressional correlations among HERV loci.

F) Association between the mean HERV expression level and the mean DNA methylation level (beta value) of the CpG sites that are on or proximal (<1 kilo base pairs (kb)) to the expressed HERVs. The result for BLCA is shown, and data for tumors and adjacent normal tissues are included. The dots are colored according to the sample type (i.e., normal tissue or respective tumor subtypes).

G) Associations of HERV expression levels and DNA methylation levels in respective types of cancers. The results for the HERV-proximal (<1 kb) CpG sites and all CpG sites are shown.

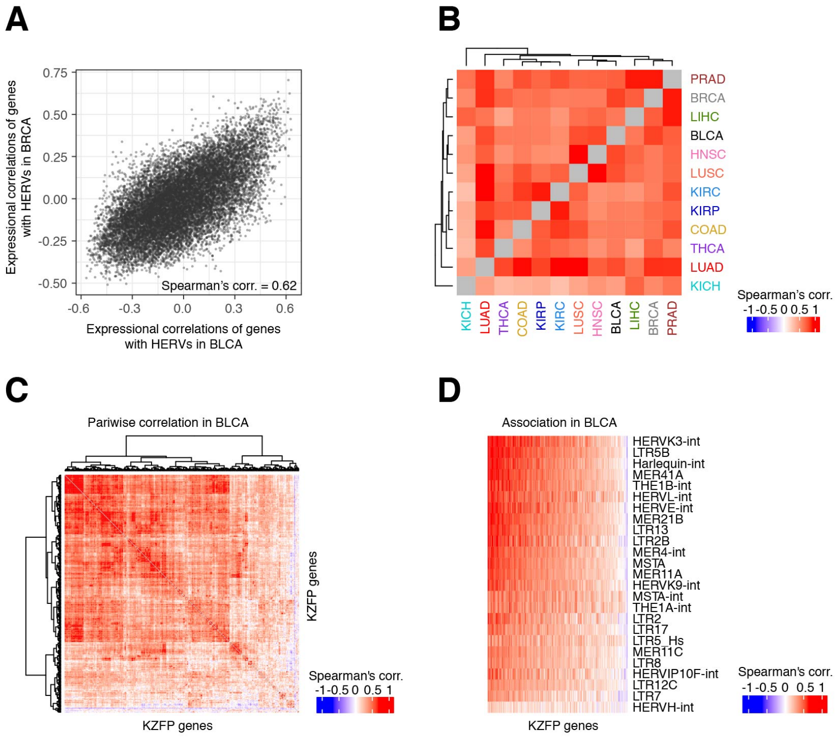


**Fig. S2 Transcriptome signatures associated with global HERV activation in tumors.**

A) Similarity of the gene expression changes upon HERV activation between the BLCA and BRCA tumor datasets. In the respective datasets, Spearman's correlation of the expression of each gene with the total expression of HERVs was calculated. Subsequently, those scores were compared between the BLCA and BRCA tumor datasets.

B) Pairwise similarities of the gene expression changes upon HERV activation among 12 cancer types.

C) Pairwise expression correlations of respective KZFP genes in BLCA tumors. The results for the 300 most highly expressed KZFPs are shown.

D) Pairwise expression correlations between HERV groups and KZFP genes in BLCA tumors. The results for the 25 most highly expressed HERVs and 300 KZFPs are shown.

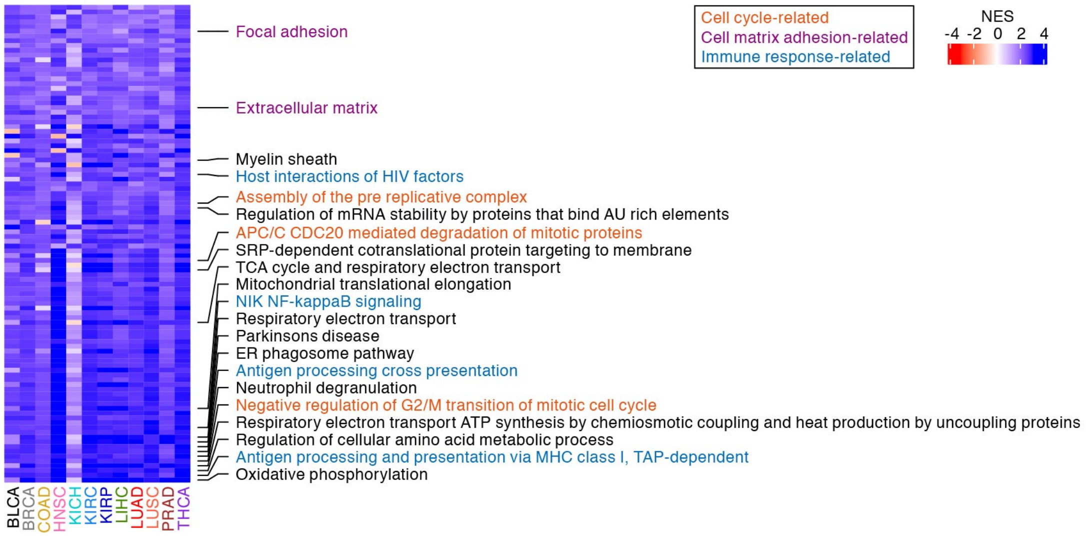


**Fig. S3 Results of GSEA summarizing genes whose expression levels were correlated with the overall expression levels of KZFP genes in the TCGA dataset.** As the overall expression level of KZFP genes, the GSVA score[46] of KZFP genes was used. The high-scored 100 gene sets with negative correlation are shown. Of these, the top 10 gene sets and the gene sets indicated in **Fig. 1F** are annotated.

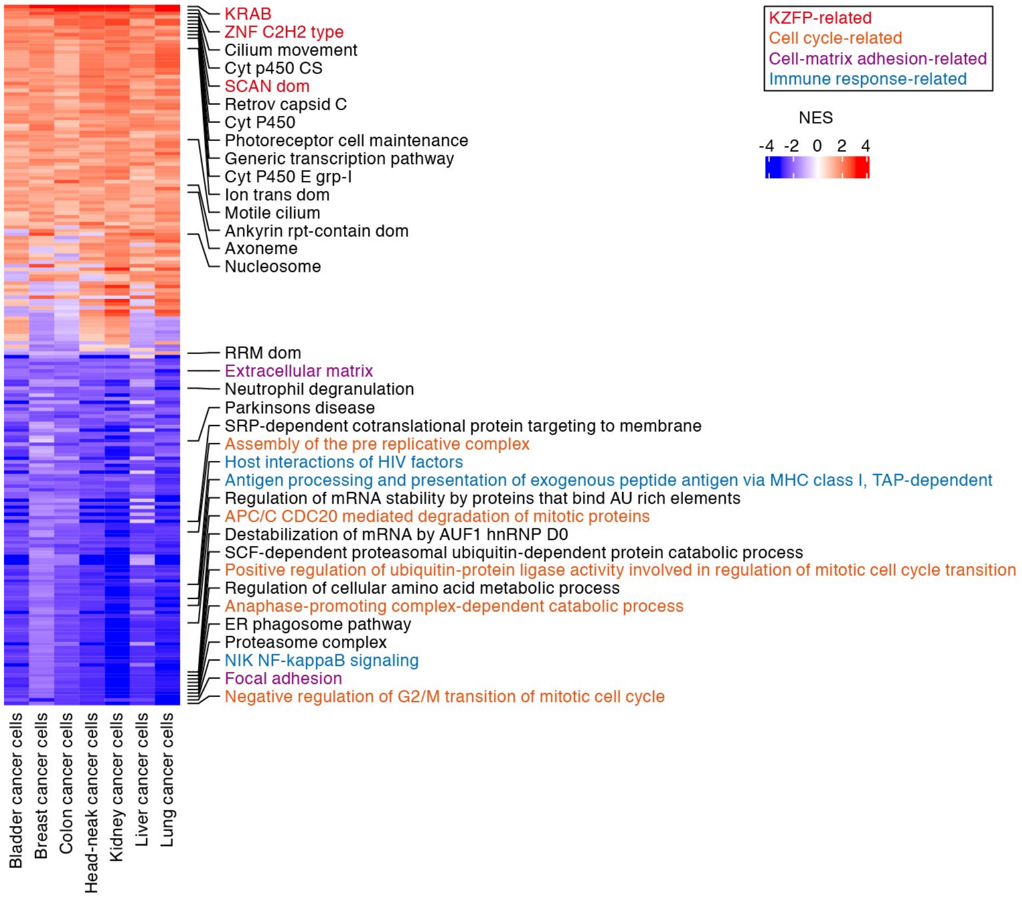


**Fig. S4 Results of GSEA summarizing genes whose expression levels were correlated with the global expression levels of HERVs in the CCLE dataset.** The high-scored 100 gene sets with positive or negative correlation are shown. Of these, the top 10 gene sets and the gene sets indicated in **Fig. 1F** are annotated.

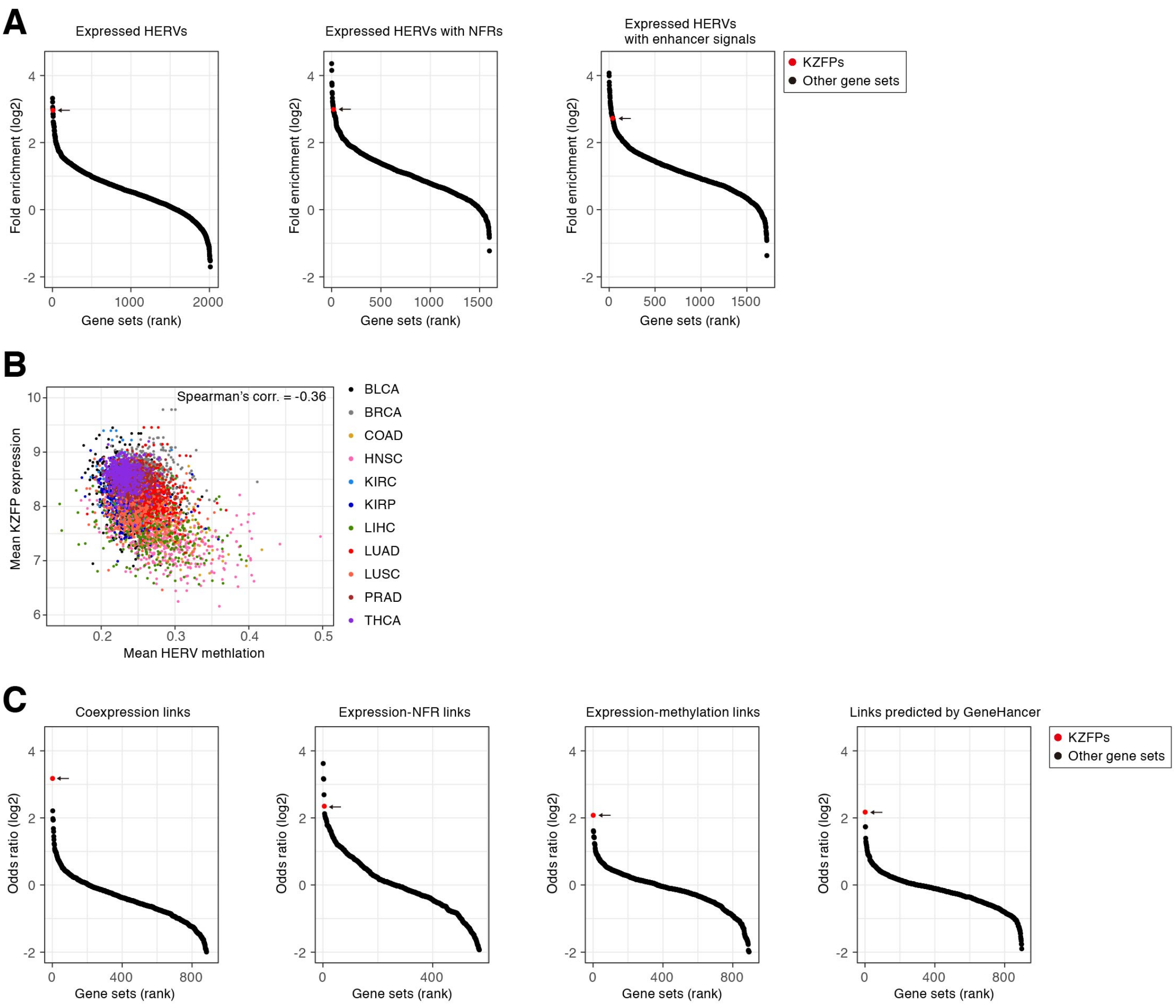


**Fig. S5 Transcriptional regulatory activity of the expressed HERVs in the vicinity of KZFP genes.**

A) Gene Ontology (GO) enrichment analyses to identify sets of genes that are preferentially present in the vicinity (<10 kb) of the expressed HERVs. The gene sets are ranked according to the fold enrichment scores, and the gene set "KZFP family" is highlighted. Only gene sets with ≥5 hits of HERVs are shown.

B) Association between the mean methylation level of CpG sites that are on or proximal (<1 kb) to the expressed HERVs in the vicinity (<50 kb) of KZFP genes and the mean expression levels of those genes.

C) GO enrichment analyses to identify sets of genes that are preferentially related to the expressed HERVs in the predicted gene regulatory network. Only gene sets with ≥1 hit of HERVs are shown.

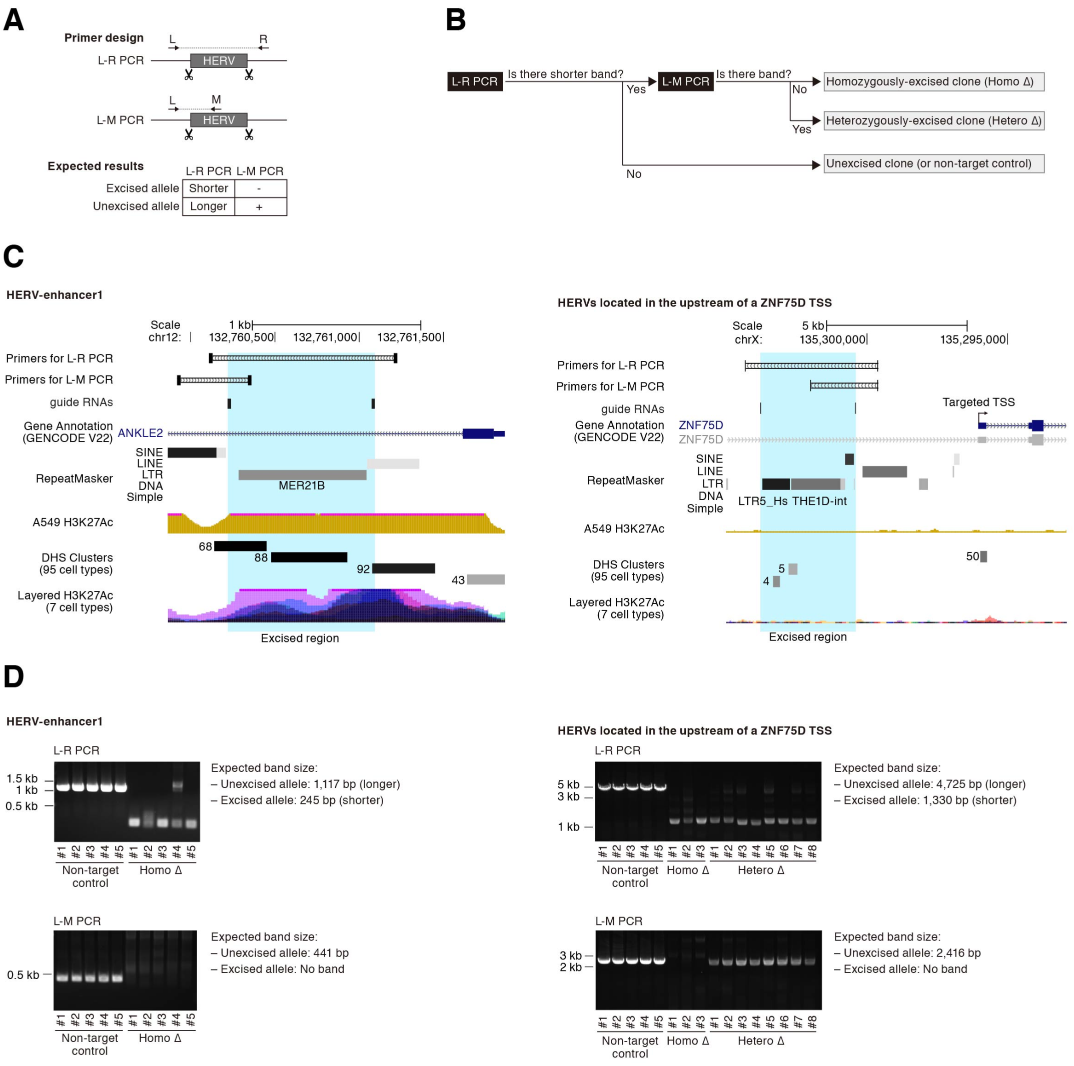


**Fig. S6 Establishment of HERV-excised A549 cell clones using CRISPR-Cas9 system.**

A) Design of PCR primers to check HERV excision. Two types of PCRs were designed: L-R PCR and L-M PCR. In L-R PCR, shorter and longer bands are amplified from HERV-excised and -unexcised alleles, respectively. In L-M PCR, a single band is amplified from HERV-unexcised allele (but not from HERV-excised allele).

B) Scheme to screen HERV-excised cell clones using L-R and L-M PCRs.

C) UCSC genome browser views of the target HERVs for excision experiments. The views for HERV-enhancer1 (left) and HERVs located in the upstream of a *ZNF75D* TSS (right) are shown.

D) PCR results of HERV-excised clones or non-target control clones. Results for HERV-enhancer1 (left) and HERVs located in the upstream of a *ZNF75D* TSS (right) are shown.

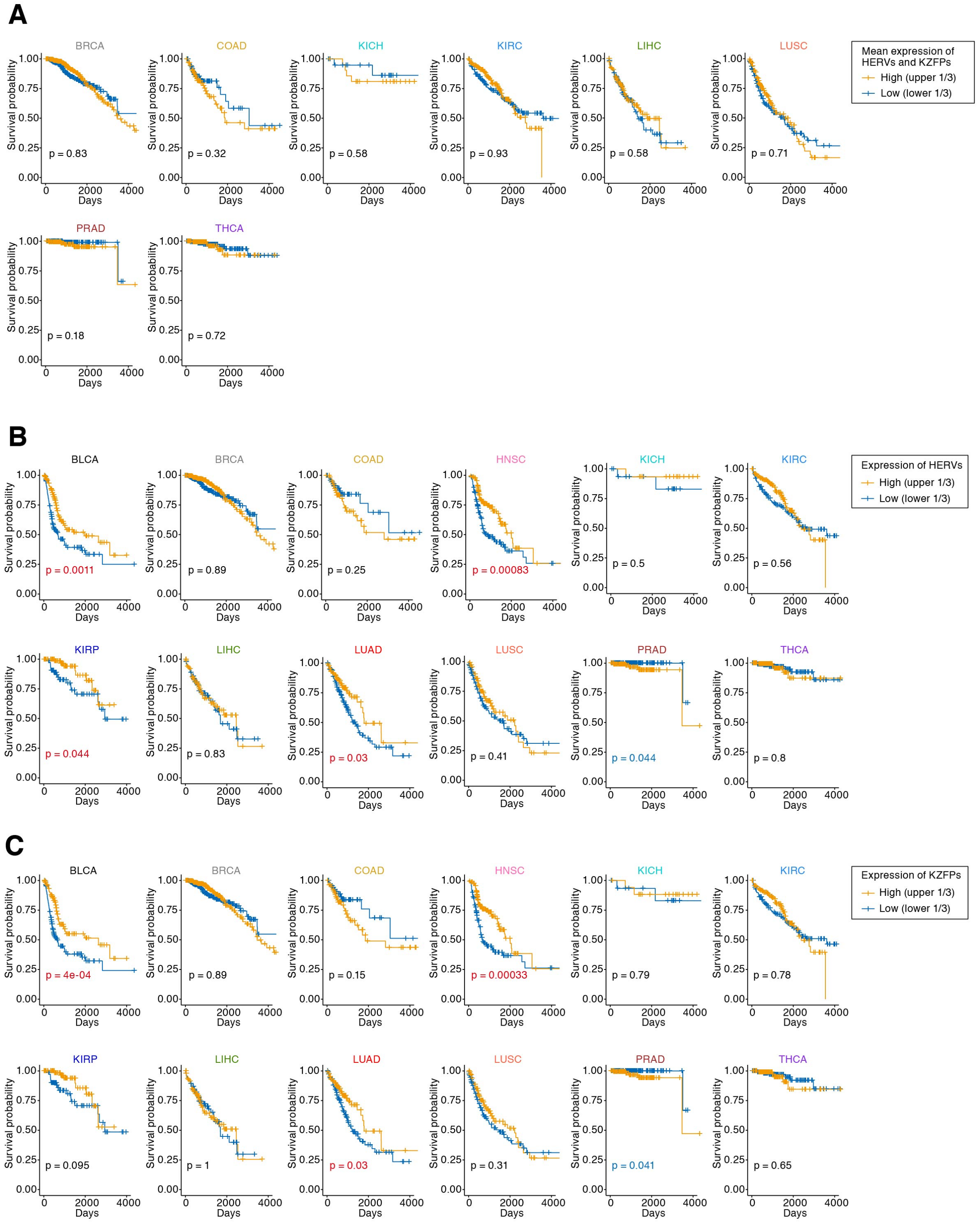


**Fig. S7 Kaplan–Meier survival plots of cancer patients with high or low expression levels of HERVs and KZFPs.**

A) Cancer patients were stratified according to the mean value of gene set-wise expression scores (GSVA scores[46]) between KZFPs and HERVs. The results for

BRCA, COAD, KICH, KIRC, LIHC, LUSC, PRAD, and THCA tumors are shown (results for the others are shown in **Fig. 3B).**

B) and C) Unlike **Figs. 3B and S7A**, patients were stratified according to the expression status of either HERVs (B) or KZFPs (C).

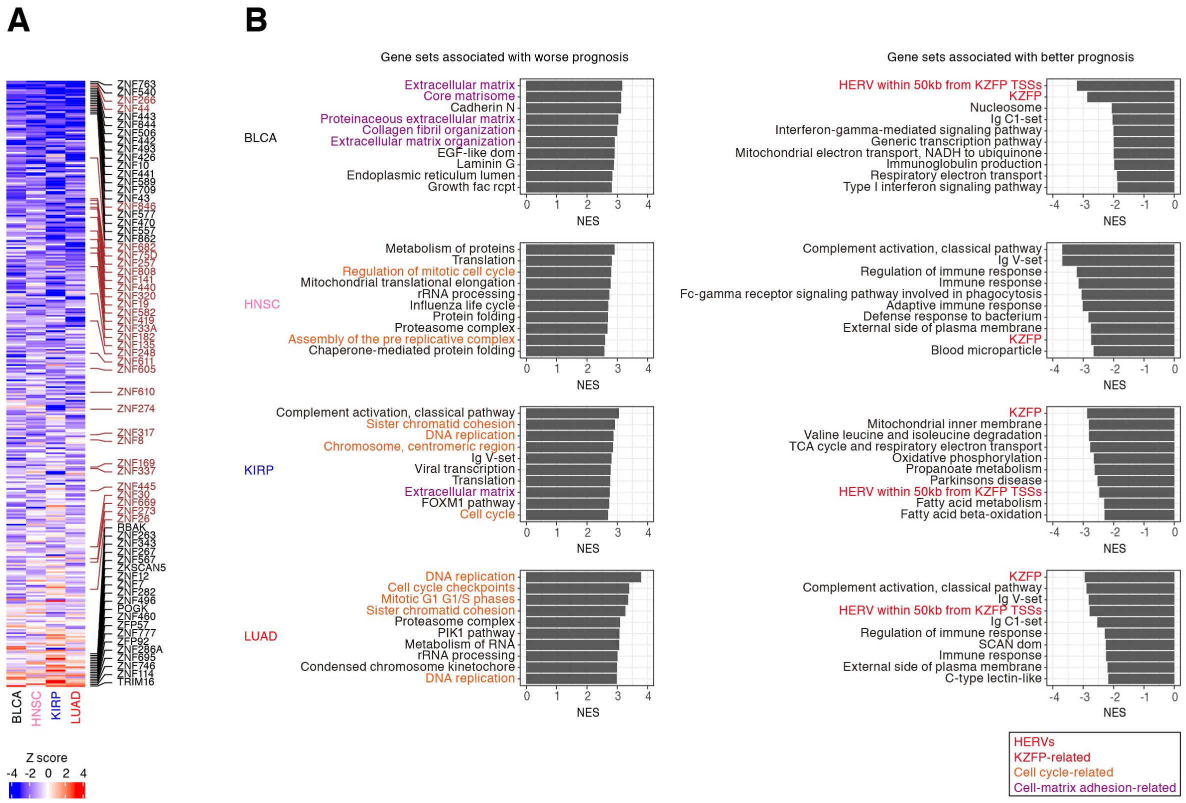


**Fig. S8 Genes associated with the prognosis of cancer patients.**

A) Associations of the expression of respective KZFP genes with cancer prognosis. The Z score in the Cox proportional hazards model is shown. The top 20 KZFP genes with respect to the association with better or worse prognoses are annotated. In addition, the KZFP genes used in the overexpression experiments (**Fig. 4**) are annotated and highlighted.

B) The high-scored gene sets from GSEA based on the Z scores in the Cox proportional hazards model. Redundant gene sets were removed from the results.

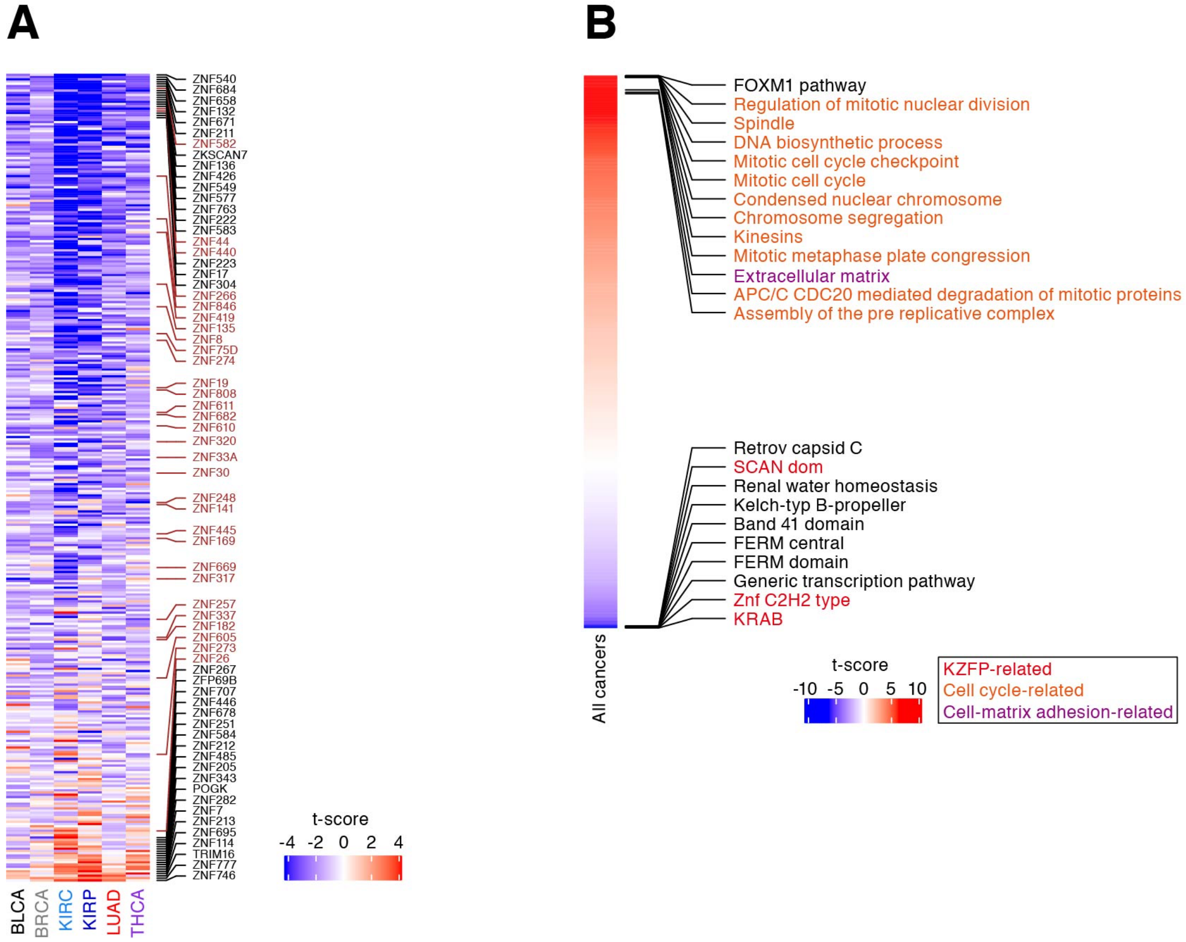


**Fig. S9 Genes associated with cancer progression.**

A) Associations of the expression of respective KZFP genes with cancer progression. The associations of the expression levels of respective genes with cancer progression were evaluated using single linear regression analysis. Positive and negative t-scores indicate the tendencies of increased and decreased expression, respectively, of the genes along with cancer progression. The top 20 KZFP genes with respect to the positive and negative associations with cancer progression are annotated. In addition, the KZFP genes used in the overexpression experiments (**Fig. 4**) are annotated and highlighted.

B) The high-scored gene sets in the linear regression analysis are shown in **Fig. 3F**. The top 10 gene sets and the gene sets indicated in **Fig. 1F** are annotated.

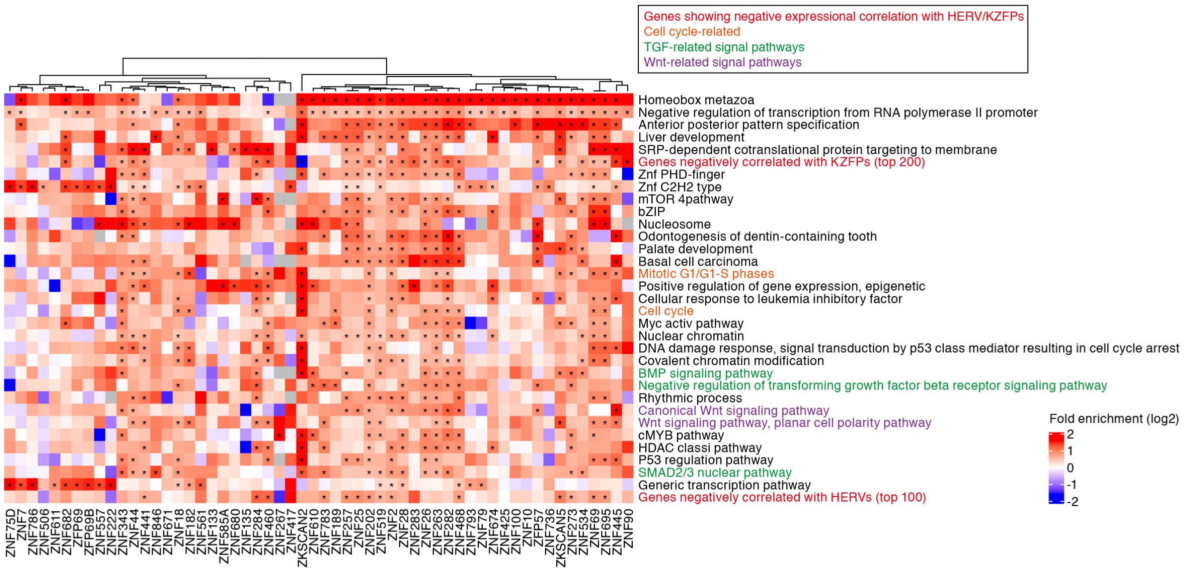


**Fig. S10 GO enrichment analysis to identify gene sets that are preferentially bound by respective KZFPs.**

This analysis was based on a publicly available ChIP-Seq dataset of KZFPs (Imbeault et al.[33]). In the heatmap, the color indicates the log2-transformed fold enrichment. An asterisk denotes a significant enrichment (fold enrichment > 1.5; FDR < 0.05). The heatmap includes 1) gene sets that were significant in >12 KZFPs, and 2) KZFPs in which ≥1 gene sets above were significant.

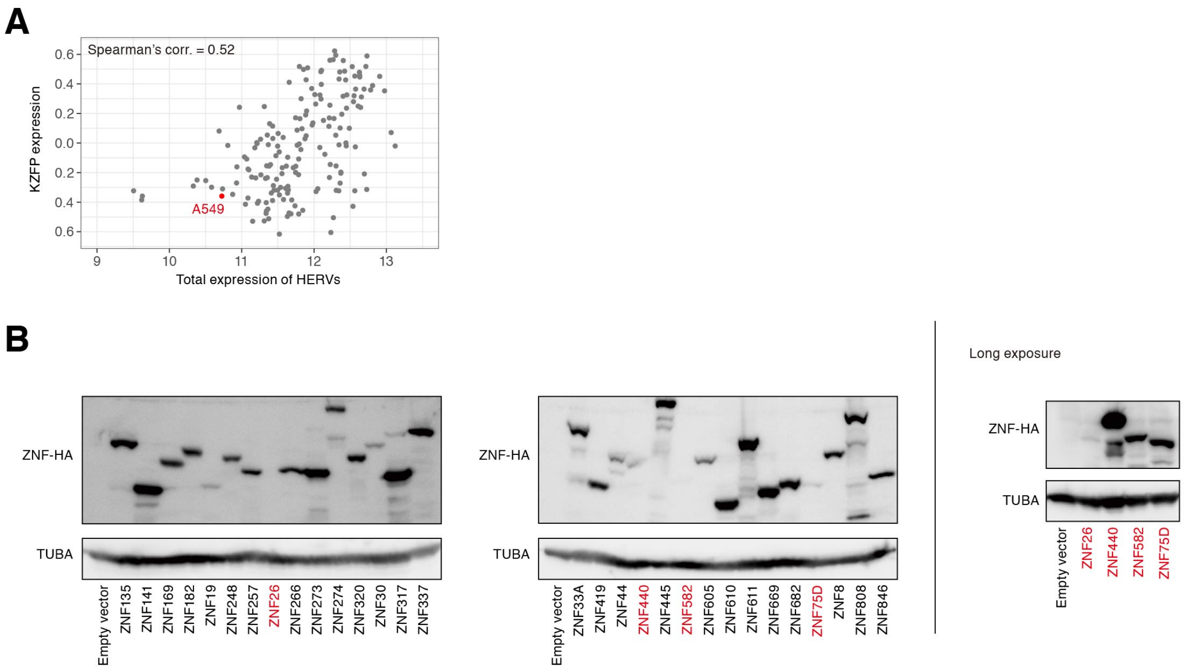


**Fig. S11 Establishment of a panel of A549/KZFP cells.**

A) Expression levels of HERVs and KZFPs in lung cancer cells in the CCLE dataset. The X-axis indicates the total expression levels of HERVs (log2-transformed CPM), and the Y-axis indicates the overall expression levels of KZFP genes (GSVA score). Dots corresponding to A549 cells are highlighted.

B) Western blotting to confirm the exogenous expression of KZFP proteins in A549/KZFP cells using an anti-HA antibody. Since the target bands in several A549/KZFP cells (indicated as red) were relatively faint (in the left panel), the results of western blotting with long exposure are also shown for these cells (in the right panel).

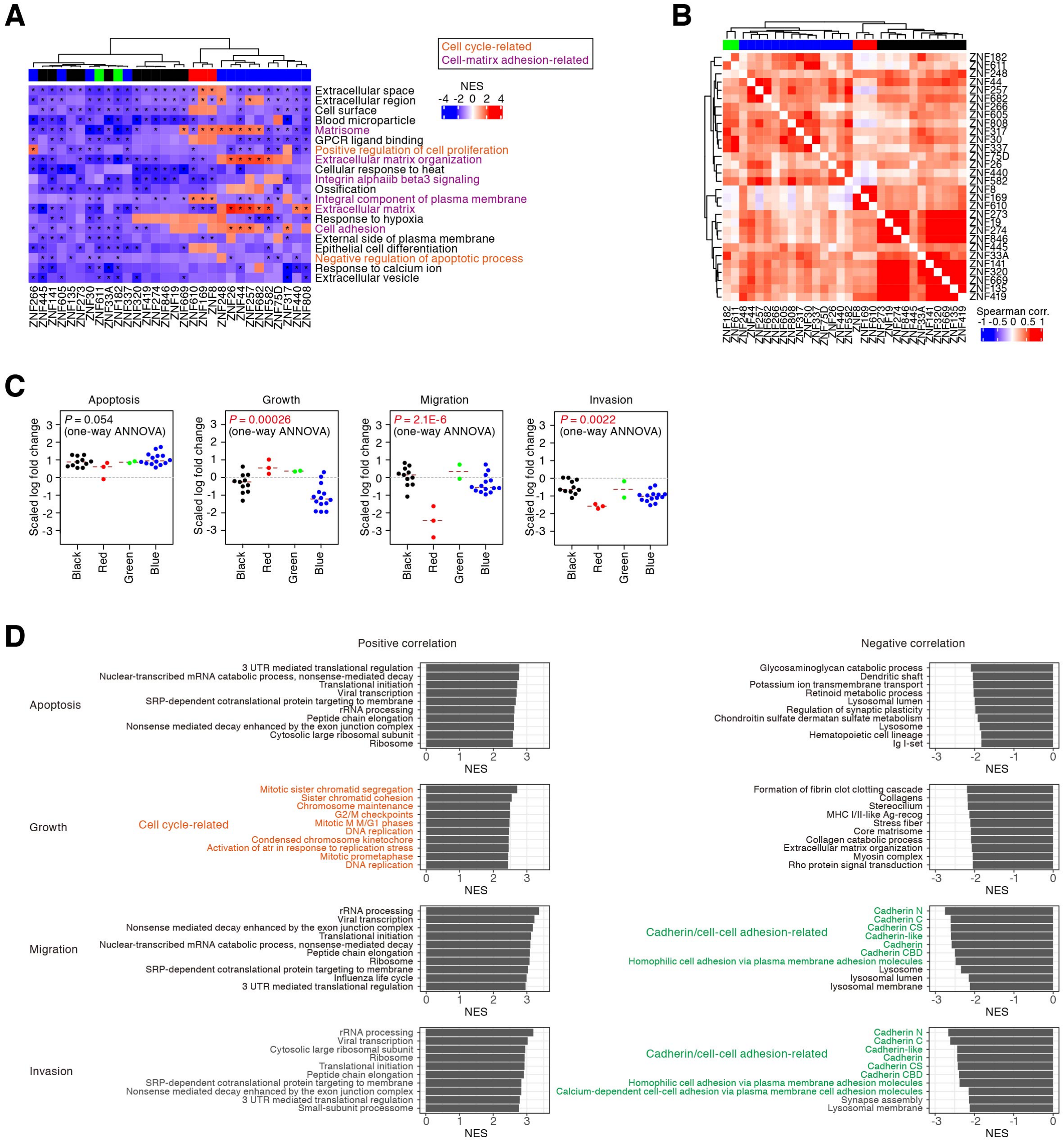


**Fig. S12 Phenotypic and gene expression changes caused by the overexpression of KZFPs in lung adenocarcinoma cells.**

A) Results of GSEA summarizing genes whose expression levels were commonly downregulated in A549/KZFP cells. An asterisk denotes a significant downregulation of the gene set in certain A549/KZFP cells. The top 20 gene sets are shown. Redundant gene sets were removed from the results. Clusters indicated in the upper side of the heatmap are the same as those in **Fig. 4E**.

B) Pairwise similarities of the gene expression alterations in A549/KZFP cells. Spearman's correlations of the fold changes of gene expression were calculated among the A549/KZFP cells. Clusters indicated in the upper side of the heatmap are the same as those in **Fig. 4E**.

C) Phenotypic differences of the cells among the gene expression-based clusters.

D) Genes whose expression levels were associated with the measured phenotypes in A549/KZFP cells. For each phenotype, Spearman's correlations of the expression levels of respective genes and the phenotype score were calculated, and GSEA was subsequently performed according to those correlation scores. Regarding the positive and negative correlations, the results for the top 10 gene sets are shown.

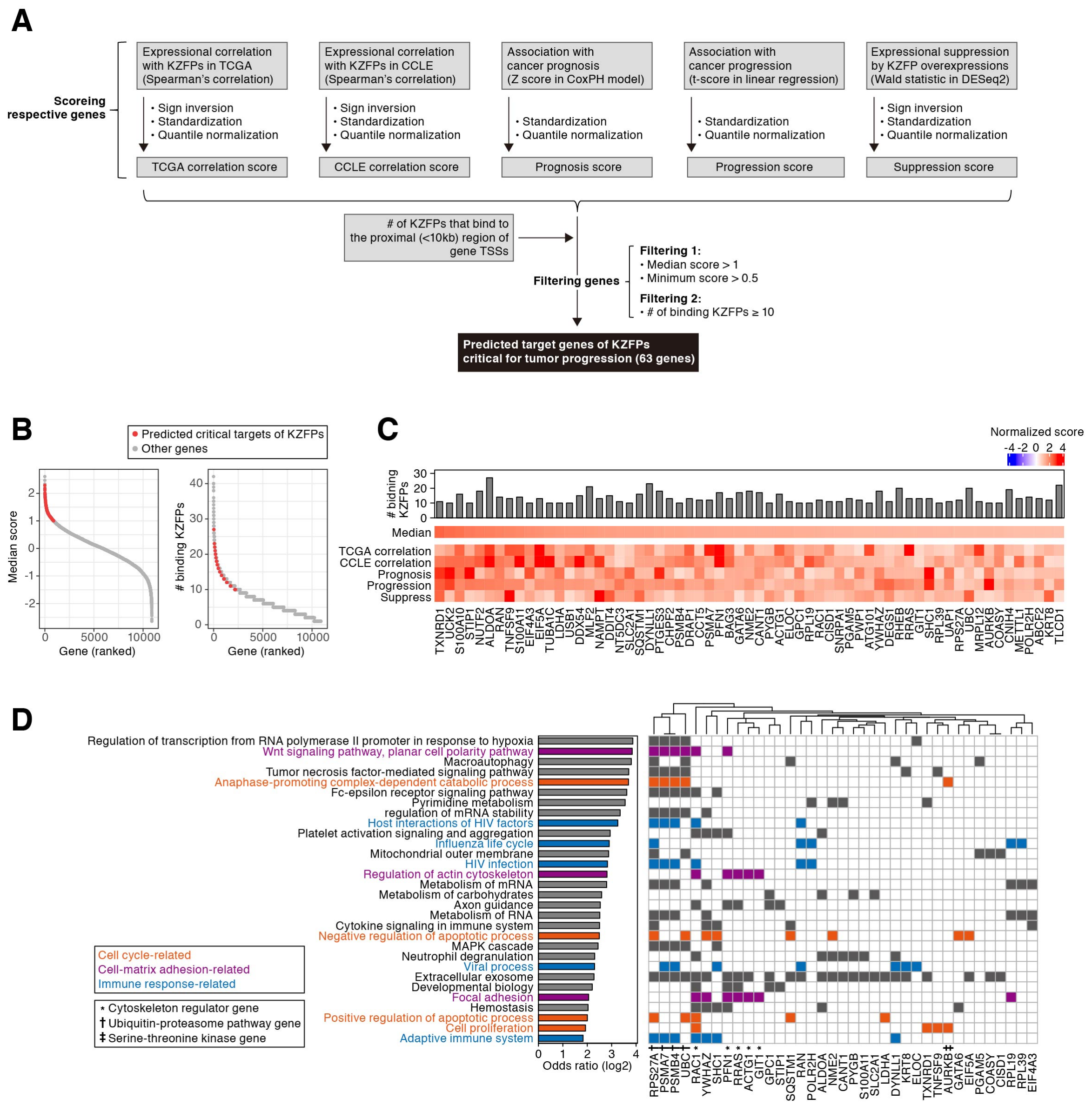


**Fig. S13 Identification of KZFP target genes that are likely to be critical for cancer progression.**

A) Scheme for extracting the possible target genes of KZFPs critical for tumor progression. For each gene, the 1) TCGA correlation score, 2) CCLE correlation score, 3) prognosis score, 4) progression score, and 5) suppression score were calculated. Genes with >1 median score and >0.5 minimum score were extracted. Subsequently, genes targeted by ≥10 KZFPs were extracted. Finally, 63 genes were extracted. See the **"Scoring system of genes for predicting the targets of KZFPs critical for cancer progression"** section in **Materials and Methods**.

B) Distribution of the median score (left) and number of binding KZFPs (right) of respective genes.

C) The number of binding KZFPs (upper), the median score (middle) and respective scores (lower) of the 63 genes.

D) GO enrichment analysis summarizing the possible target genes of KZFPs critical for cancer progression. Of the significant gene sets (i.e., FDR < 0.1; number of hits ≥ 5), the top 30 gene sets according to the odds ratios are shown. Log2-transformed odds ratios (left; bar chart) and gene memberships (right; heatmap) are indicated. Cytoskeleton regulator genes (*), ubiquitin-proteasome pathway genes (†), and serine-threonine kinase genes (‡) are annotated under the heatmap.

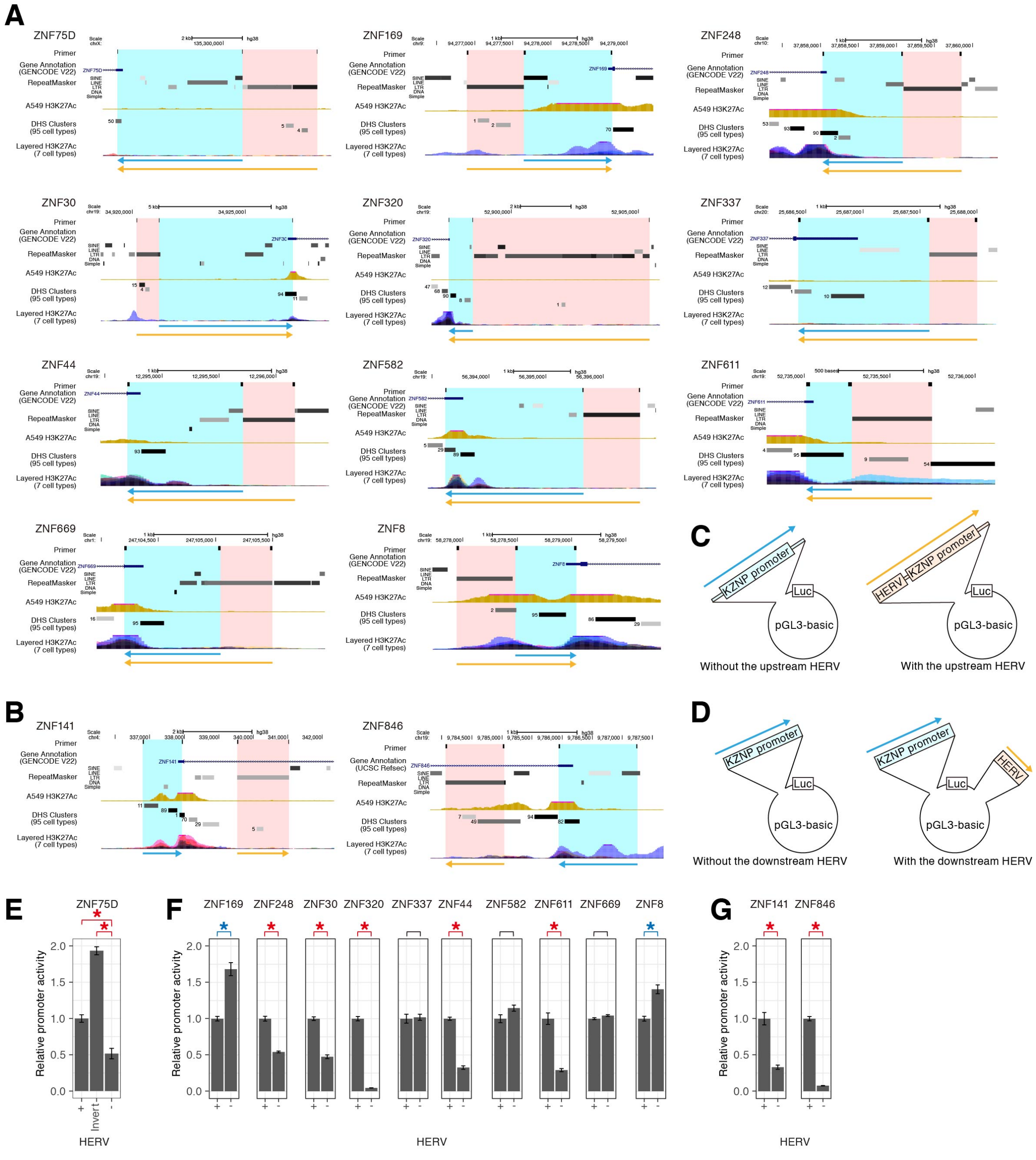


**Fig. S14 Luciferase reporter assay to assess the effects of the surrounding HERVs on the promoter activities of KZFP genes.**

A) and B) UCSC genome browser views of the target HERVs and KZFP promoters for the luciferase reporter assay. The panel for a HERV in the upstream region of the KZFP promoter is shown in A), while that for a HERV in the downstream region is shown in B). The genomic region inserted into the reporter plasmid is indicated by the arrow. In A), the orange or blue arrows indicate genomic fragments with or without HERV sequence, respectively. In B), the orange or blue arrows indicate HERV or KZFP promoter, respectively.

C) and D) Schematics of the reporter plasmids. Schematic for a HERV in the upstream region of the KZFP promoter is shown in C), while that for a HERV in the downstream region is shown in D).

E) Assessment of the directional effect of the HERVs on the promoter activity of *ZNF75D*.

F) Assessment of the effect of the upstream HERVs on the promoter activity of the KZFP gene.

G) Assessment of the effect of the downstream HERVs on the promoter activity of the KZFP gene.
